# Volume conductor models for magnetospinography

**DOI:** 10.1101/2024.11.04.621905

**Authors:** George C. O’Neill, Meaghan E. Spedden, Maike Schmidt, Stephanie Mellor, Matti Stenroos, Gareth R. Barnes

## Abstract

The recent development of small, wearable, magnetic field sensors allow for the investigation of biomagnetic fields with a flexibility previously unavailable. We carry out forward computations to describe how current flow in the spinal cord and thorax gives rise to measurable magnetic fields outside the torso. We compare various open-access volume conductor models, in order to select the most parsimonious and accurate descriptor of the magnetic fields due to source current in the spinal cord.

We find that fields produced due to current flow along the superior-inferior axis of the cord are relatively insensitive to the choice of volume conductor model. However, fields produced by current flow in predominantly left-right or anterior-posterior direction are significantly attenuated by the presence of bone in the forward model. Furthermore, volume conductors with bone demonstrate larger differences in field topographies for nearby sources compared to bone-free models. These findings suggest that precise modelling of spinal cord location and surrounding vertebrae will be important a-priori knowledge going forward.

## 1) Introduction

The spinal cord and its associated activity in sensorimotor networks are of interest to both basic neuroscience and clinical research. From a neuroscience perspective, understanding its integrative role in combining ascending afferent activity from the body to fine-tune descending motor control is an area of interest for researchers^1 3^. Whilst in the clinic it is desirable to non-invasively identify the location and severity of spinal cord injury^4^ and track any potential recovery from treatment approaches^5,6^.

One approach to measure activity originating from the spinal cord which is being developed is magnetospinography (MSG), where the activity is measured non-invasively from the magnetic fields generated by currents due to the neuronal activity. The potential advantages of using the magnetic field to measure spinal cord activity are twofold. First, the magnetic fields can be sensed without direct contact to the subject, avoiding the need for electrodes placed on the skin or invasively in the epidural space^7,8^. Second, magnetic fields originating from neural activity are less affected by the poorly conducting bone than the corresponding electric potential, and overall the uncertainty of conductivity values of tissues affect magnetic signals less than electric signals, making modelling easier^9,10^. Therefore, if we were to localise where in the spinal cord these signals originated from, we should have an improved spatial resolution compared to the non-invasive measures based on electric potential (e.g. electrospinography^11,12^).

Measuring magnetic fields from the spinal cord is a challenge due to its depth (∼50 mm) from the skin surface and the relative low amplitude of the expected current moment (∼10 nAm^13^). Quantum sensors with femtoTesla level sensitivity are required for this task and so initial research into MSG recordings have been performed using bespoke Superconducting Quantum Interference Devices (SQUIDs) systems^13 17^. These systems generate high quality data which can localise the spatiotemporal properties of the neuronal activity, but the bulky cryogenic support infrastructure means it can only image a limited portion of the spinal cord per experiment^15^. Recent advances in cryogen-free sensors (such as optically pumped magnetometers; OPMs) allow us similar sensitivity but with sensors the size of a 2×4 LEGO brick which can be flexibly placed anywhere near or on the back, allowing for the possibility of covering large areas of the back with OPMs and any other location relevant for measuring the peripheral nervous system. To this end, the first experiments with OPMs to measure spinal activity are coming online^18,19^.

Irrespective of acquisition method, to fully leverage the source localisation abilities promised by MSG, we need to ensure we are approaching the forward problem (modelling how a known source current distribution is represented at the sensor-level) and inverse problem (estimating the current distribution from set of sensor-level observations) in a manner appropriate for the spinal cord. Previous source analyses of MSG data have used simple volume conductor models to solve the forward problem, such as assuming an infinite homogenous medium^17^ or a basic approximation of the torso shape^18^.

This paper compares and contrasts a set of existing volume conductor modelling approaches from the magnetoencephalography (MEG) and magnetocardiography (MCG) literature, implemented in academic software toolboxes on a theoretical OP-MSG setup, to investigate the similarities and differences between them in the context of magnetospinography. Similar to previous encephalographic conductive model comparison studies^10,20,21^, we test and compare increasingly complex volume conductors to understand the benefits additional modelling provides. Based on the results presented here, we make some recommendations on selecting an appropriate volume conductor for MSG.

## 2) Methods

### 2.1 Model geometry and source space

Our simulations are based on the anatomy and posture of a participant who undertook a previous OP-MSG study^18^. **A scan of the particpant’s head and torso whilst they were seated was generated using** an infra-red structural camera (Occipital Inc, Boulder, CO). The geometry is shown in Figure 1A. To generate a basic boundary of the torso, we modified the thorax mesh provided in ECGsim^22^. First, we upsampled the ECGsim thorax, cardiac blood and lung meshes. Then, we included an abdomen and neck to form a new torso mesh and registered the mesh to the participant scan with a two-step process: an initial 7 degrees of freedom (translation, rotation and global scaling) fit using three fiducial locations, the left and right acromion, and L5 point of the spine was followed by a constrained iterative closest point fit to generate the full 12 degrees of freedom affine transformation. An cartoon example of this fit can be seen in Figure 1E. This transformation was also applied to the heart and lung meshes from ECGsim.

**Figure 1.**
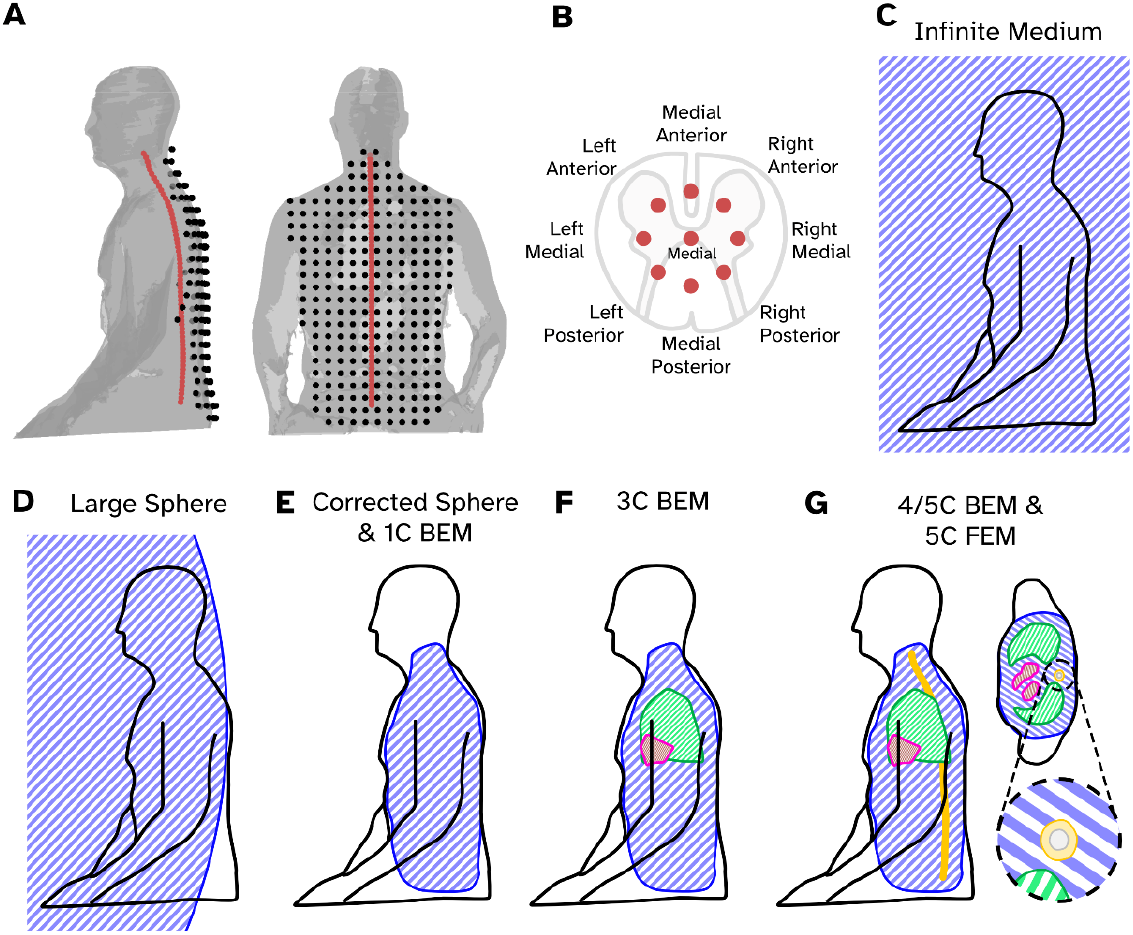
Setup of the source, sensor and volume conductor models for the simulations. A) The sensor (black dots) and medial source locations (red dots) relative to a structural scan of the example participant (grey surface). B) The locations of 9 modelled source locations for a transverse “slice” of spinal cord, a cartoon slice of spinal cord has been superimposed to illustrate where these sources are sat. C-G) Cartoon diagrams showing how the main conductivity boundary for each tested forward model (blue shaded area) is positioned. In addition to the main volume, sub domains representing the lungs (green), heart (red), spinal cord (grey) and vertebrae (yellow) and depicted. Panel G also includes a transverse slice through the chest to show the locations of the conductivity boundaries in more detail.

With the torso mesh fitted, we approximated the curvature of the spinal cord using a 5^th^ order polynomial function and a source space was generated 50 mm deep with elementary sources placed 10 mm apart (see red dots in Figure 1A). For every elementary source (labelled as the medial source in Figure 1B), 8 other source locations, were placed to form a ring with 4 mm radius around the medial source (Figure 1B). This ring of sources was allowed to rotate along the plane of the ring to follow the curvature of the spine. Having generated a source space for the spinal cord, we also generated two additional meshes to represent the spinal cord conductive volume which wrapped the spinal cord with an 8 mm radius from the medial sources, and a second larger mesh representing the vertebrae with a radius of 16 mm from the medial sources. (see Figure 1G for an example). Table 1 provides the details of the meshes used for this study. We tested whether the densities of the meshes provided in Table 1 were sufficient for BEM analysis, these results can be found in the Supplementary Material.

**Table 1.** The meshes used to generate the conductive models tested.

| Name | Nodes | Median Node Separation (mm) | Elements | Conductivity (Siemens/m) |
| --- | --- | --- | --- | --- |
| Torso | 3160 | 15.8 | 6316 (faces) | 0.23 |
| Heart | 1052 | 6.4 | 2906 (faces) | 0.62 |
| Lungs | 914 | 15.8 | 1820 (faces) | 0.05 |
| Spinal Cord | 5161 | 2.7 | 10318 (faces) | 0.33 |
| Bone | 4141 | 5.1 | 8278 (faces) | 0.007 |
| FEM combined | 103131 | 7.0 (across entire volume)<br>2.0 (within spinal cord domain) | 711030 (tetrahedra) | Element specific (see above) |

**Table 2.**
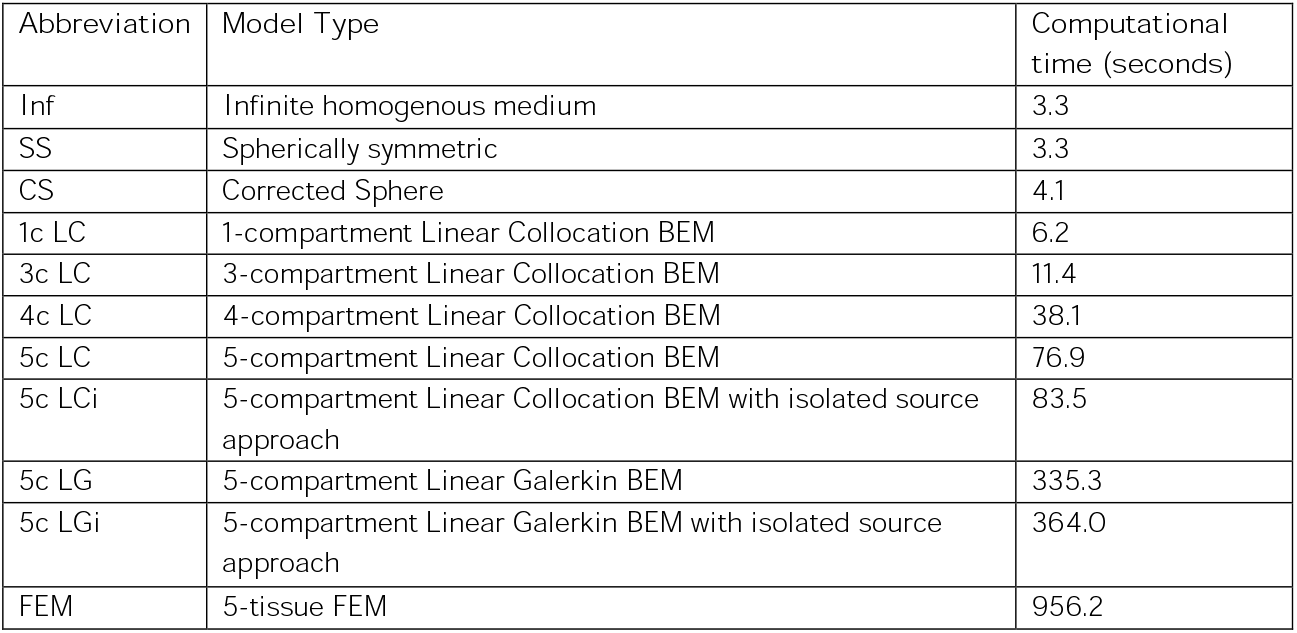
A list of the 11 volume conductor models tested, with their abbreviations used in the results, and the time it took to compute the field topographies using a workstation with an Intel i9-12900 CPU, 64 GB of RAM running MATLAB 2023a on Windows 11.

Sensor locations were generated using a 2D grid of sensors (spaced 30 mm apart) which were then ray-cast onto the structural scan of the participant, keeping only the sensors which would also ray-cast with the fitted torso mesh. Sensors were placed 10 mm from the structural scan surface, which is approximately the distance the sensors are from the scalp in OP-MEG experiments^23,24^. This generated 250 sensors (black dots in Figure 1A). The sensors were oriented such that their primary axis was along sagittal axis of the body, and tangential fields were measured along the frontal and transverse axes for a total of 750 channels. Triaxial measurements were chosen as both SQUID-based and OPM-based magnetospinographs already offer multi-axis recordings ^13,18^. The sensors were modelled point-like, i.e. the extent of the sensor was not accounted for in the model.

### 2.2 Volume conductor models

Bioelectromagnetic fields are commonly modelled in terms of quasi-**static approximation of Maxwell’s** equations in an ohmic conductor^25,26^. In the quasi-magnetostatic approximation, current is assumed to flow in closed loops, i.e. total current density 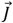 is divergence-free, 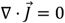. The total current 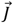 is divided into the primary source current 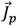 that represents the source activity at the macroscopic scale, and volume current 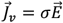 that is driven in the conductor of conductivity *σ* by the electric field 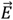 that is created by charge density associated with the divergence of 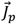. Under quasi-electrostatic approximation, 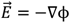, yielding, 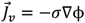. From 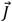, one can evaluate the magnetic field 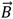 at sensor position 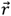 using the Biot-Savart formula. It is practical to separate the field to 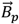 and 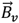, corresponding to sources 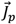 and 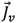. We discretise 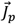 into a set of current dipoles 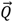, so it suffices to write the formula of 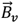 for a current dipole 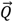 at position 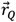 only:

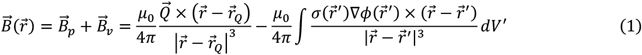

#### 2.2.1 Simplified volume conductor models

The simplest volume conductor model is an infinite homogenous conductor (Figure 1C). in this case the volume current 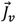 does not contribute to the magnetic field, thus 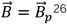.

If *σ* is spherically symmetric and finite, 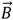 outside the conductor is due to an internal dipole has a closed-form solution26. The field does not depend on the radial profile or conductivity or *σ*, but only its origin. The sphere model has been popular in MEG source analysis, performing well in motor and somatosensory cortices21. We set the origin approximately 1000 mm in front of the participant so that **the sphere’s surface approximated the curvature of the spine (which we denote as the Large Sphere;** Figure 1D).

The next level of realism is a homogenous finite volume conductor of arbitrary shape. Such a model can, avoiding element analysis, be solved by perturbing a sphere model using the a harmonic basis set to approximate the shape of the conductor^27^. This approach, called the corrected sphere (or single shell) model is commonly used in MEG and performs as well as a corresponding single compartment model formulated using boundary elements^21^. Here, we placed the origin of the sphere within the torso boundary and fitted spherical harmonic gradients of the torso boundary up to an order of 𝓁=10 to form our perturbator. The conductive boundary used to fit the spherical harmonic gradients can be found in Figure 1E.

#### 2.2.2 Element Models

To solve 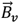 in a realistic geometry, one needs to solve the electric field 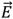 or potential ϕ. For electric potential ϕ the relations above yield

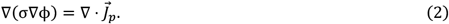

When *σ* is piecewise homogenous, the 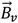 term of Eqn. 1 can be expressed as

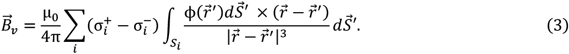

Where 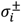 are the conductivities outside and inside boundary ***S***_***i***_. To solve 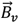. We need to know ϕ only at boundaries ***S***_***i***_. The boundary potential can be solved by converting Eqn. 2 into a surface integral form, discretising the boundary surfaces and potentials. That system is solved using the boundary element method (BEM).

We tested 5 different variants of conductivity distributions, primarily by increasing the number of different tissue types. The 1c BEM (Figure 1E) incorporates just the torso volume; it should be roughly equivalent to the corrected sphere model. The 3c BEM also included the heart and lung meshes (Figure 1F); the 4c BEM included the spinal cord mesh; and the 5c BEM also included the bone mesh (Figure 1G). Having sources close to strong jumps of conductivity may lead to numerical issues that are alleviated by the Isolated Source Approach^28,29^. We also tested the effect of isolation at the inner boundary of the spinal cord. The conductivities of each of the tissue types are as follows: 0.33 Sm^-1^ for the spinal cord, 0.007 Sm^-1^ for the bone, 0.62 Sm^-1^ for the heart, 0.05 Sm^-1^ for the lungs and 0.23 Sm^-1^ for the torso. These conductivities are based on measurements from previous studies^30,31^, though we note that reported measures of the conductivity of tissue can be highly inconsistent^31,32^. We applied two different approaches to solve for the potentials on the surfaces. First we used the linear collocation method via the Helsinki BEM Framework^9,33^ (https://github.com/MattiStenroos/hbf_lc_p), for all models from the 1c BEM to 5c BEM, and a linear Galerkin approach^29^ for the 5c BEM models. The four BEM solvers in the text are denoted LC (linear collocation), LG (linear Galerkin) and LCi, LGi to represent variants with the isolated source approach included.

Eqn. 2 can also be discretised using basis functions in the entire conductor volume, solved using the finite-element method (FEM). The *σ* does not need to be piecewise homogenous, nor isotropic. From the solved volume potential one can extract 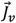 and calculate the 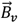 of Eqn. 1. With the FEM it is possible to make and solve very detailed volume conductor models, and is the most computationally intensive approach we test in this study. Here we took the surface meshes used in the BEM models and converted them into a single tetrahedral mesh with the ISO2MESH toolbox to prepare them for finite element analysis, with the constraint that no single tetrahedron could be any larger than 10 ml in volume. We considered the conductivity values to be homogeneous within each tetrahedron and their corresponding conductivity values were the same as for the BEM; in other words, we implemented a piecewise-homogeneous volume conductor model. The FEM was solved using the DuNeuro library^34^, in particular using the binaries compiled for the Brainstorm software suite^35^. We used the Lagrange (or continuous Galerkin) method with a restricted St. Venant source model to represent the volume currents^36^. The St. Venant model uses a weighted set of monopoles on nearby connected nodes of the mesh to approximate 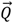, which allows us to avoid mathematical singularities^36^. Here the restricted mode implies that all monopole sources can only exist within the spinal cord domain of the mesh.

### 2.3 Model Evaluation

#### 2.3.1 Direct comparison between lead field patterns

For a given forward model at each of the candidate source locations, three dipoles oriented along the cardinal axes were generated. For a given source and orientation we compared lead fields from all models with two metrics. First, for a pair of field topography vectors 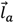 and 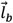, the relative error is:

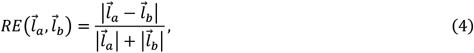

where 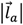 represents the L2-norm of the lead field pattern 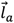. Note that this metric, in contrast to other field comparison studies^20,21^, is symmetric and non-negative, where 0 is identical and scores of 1 and higher represents dissimilarity. The limitation of the relative error measure here is that when the error is high, we cannot disambiguate whether this is due to a global gain error in the lead field or whether that the two are uncorrelated. We therefore also consider the squared correlation coefficient between 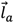 and 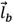:

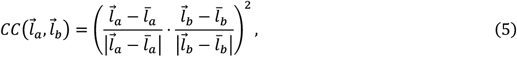

where 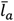 is the mean of the vector 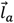.

#### 2.3.2 Decomposition of orientation sensitivity

Given the model current flow in all three cardinal directions for a given source, it is possible to determine which orientations we have sensitivity for a given volume conductor. For a triplet of field topographies 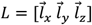, where 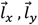 and 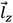 are column vectors representing the field topographies of current flow in three orthogonal orientations from the same point in space, we can perform a singular value decomposition (SVD) of the matrix such that

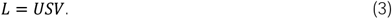

The columns of ***U*** represent the each of the 3 normalised **‘eigenfields’**, the diagonal elements of ***S*** represent the relative intensities of the eigenfields and the rows of ***V*** represent the orientation of the current flow of the corresponding eigenfields relative to the original coordinate frame.

## 3) Results

Figure 2 shows an example of a lead field topography generated with a finite element model (FEM). Equivalent plots for all the other conductive models tested can be found in the Supplementary Information. The source originates from approximately the T9 area of the spinal cord (see bottom right panel red dot). The dipole has been oriented in each of the three cardinal orientations. For each lead field we plot each axis from the triaxial OPMs we simulated, with the Y-axis here defining **the “axial” orientation (oriented normal to the** plane of the screen). X (inferior-superior) and Z (left-right) correspond to sensitivity tangential the surface of the back. The first observation is that for all three orientations of source, the sensors with the maximal sensitivity are tangentially oriented. Next, as expected the dipole oriented normal to the plane of sensors (A-P) produces the lowest amplitude fields (0.4 fT/nAm). The dipole oriented inferior to superior (I-S) has a maxima which is approximately 6 times larger than the maxima of the R-L oriented source (9.5 fT/nAm for the I-S source vs. 1.6 fT/nAm for the L-R source). On observing this result, we hypothesised this may be due to the conductivity and morphology of the bone compartment; current flow along the inferior-superior axis of the spine will be unattenuated compared to off-axis current flow. We investigate this further later in the results.

**Figure 2.**
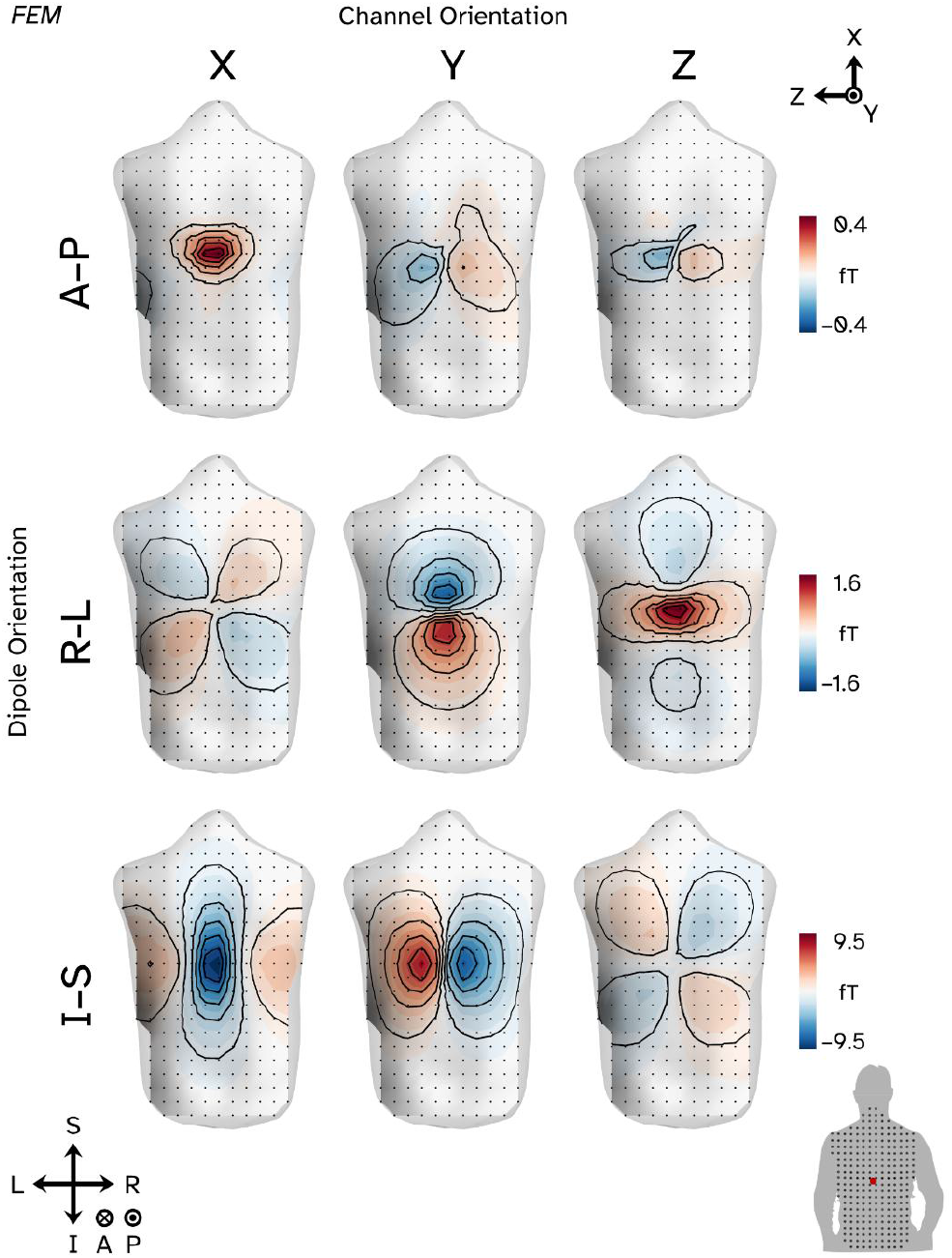
Sensor level field topographies for a 1 nAm current dipole simulated in the thoracic region of the spine (approximately T9; see red dot of bottom right panel) when the sources are oriented along the anterior/posterior (A-P) axis of the body (top row), right/left (R-L) axis (middle row) and inferior/superior (I-S) axis (bottom row). The plots are sorted into channels with a common orientation in rows. X and Z (left and right columns) represent channels oriented tangential to the surface of the back whilst Y (middle column) represents the channels oriented normal to the back. Black dots represent sensor locations and black lines represent field contour lines (separating undeciles of field strength). Similar plots for other conductive models can be found in Supplementary Information.

### 3.1 Direct comparisons between models and solutions

In Figure 3, we directly compare the field patterns produced by the 61 medial (c.f. Figures 1A-B) sources under different forward modelling assumptions to one another with our error and correlation metrics. Each element of Figure 3A is the median relative error between a pair of models when we include field topographies generated from source currents flowing in any of the three cardinal orientations. We see two distinct clusters of models with low error relative to each other. First are the conductor models which all contain bone (5c BEMs and FEM). Second are the bone-free numerical models (1c-4c BEMs), and then the three analytical solutions (infinite and spherical models) that do not cluster with anything. Focusing on the correlation between models (Figure 3D), we see that the majority of models show very high similarity to each other, with all the analytical models (1c BEM up to FEM) showing a median squared correlation of 0.95 or higher. To aid visualisation of the matrices we have plotted the errors and correlation between the FEM and all other models (Figure 3G), in other words just the final row or column of Figures 3A and 3D. We can see that relative to the FEM, the models containing bone (right of the dashed line in Fig. 3G) have a low error and high correlation, the bone-free numerical models (1c-4c BEM) have larger errors but still high correlation, finally the simplified models (Inf, SS, CS) showing the largest errors and lowest correlations.

**Figure 3.**
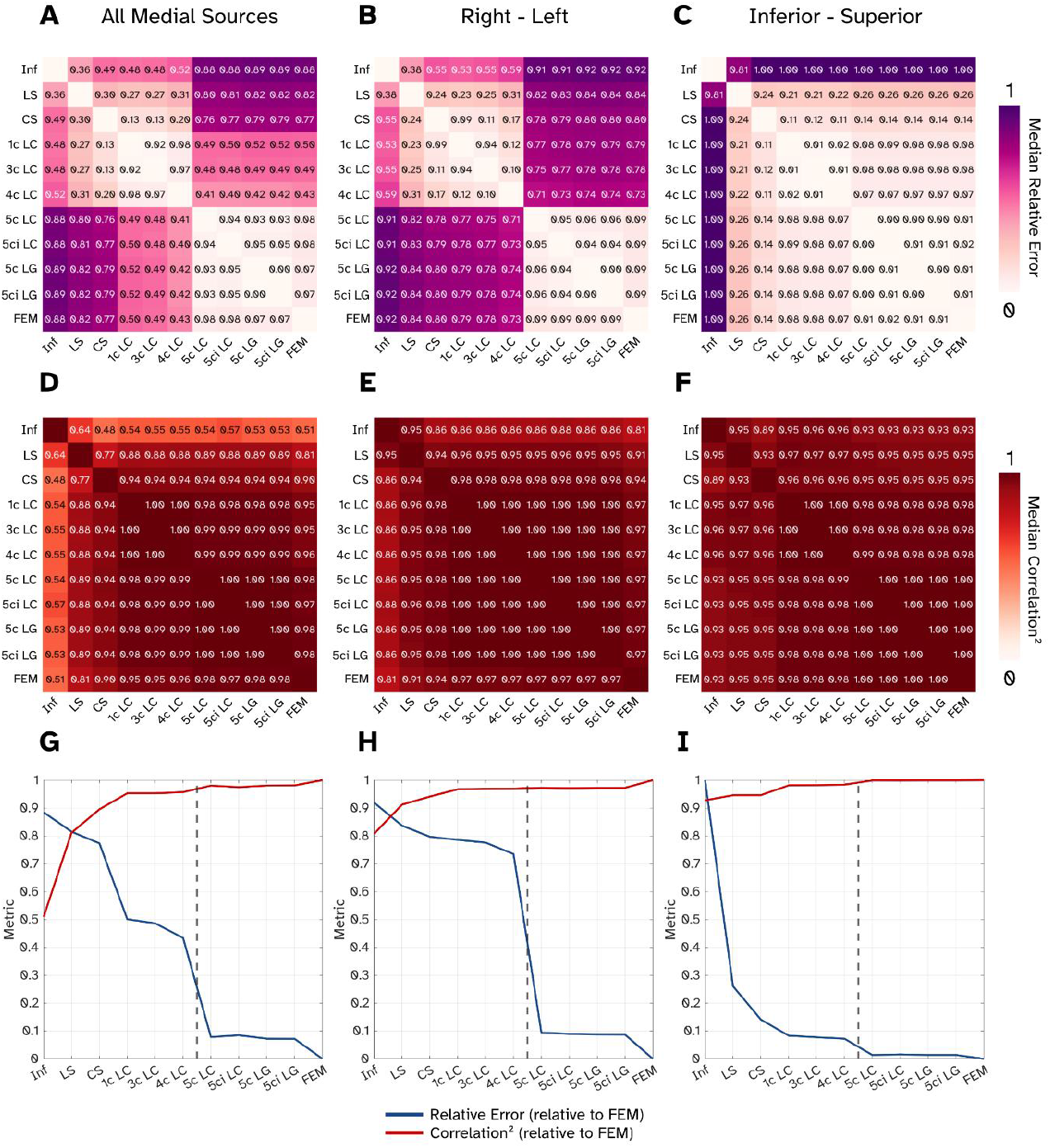
Assessment of the similarities and differences between all 11 volume conductive modelling approaches when investigating only the medial sources. **A-C)** The median relative error between pairs of models represented in matrix form. Here errors are displayed when current flow is modelled in all three cardinal orientations (A), only sources oriented right-left (B) and inferior-superior (C). **D-F)** The median correlation coefficient (squared) between pairs of models for all orientations (D), right-left (E) and inferior-superior (F). **G-I)** Both the relative and correlations plotted, but this time between the Finite Element Model (FEM) and all other models for all orientations (G) right-left (H) and inferior-superior (I). Black dashed line shows the boundary between models containing bone (right of line) and not.

To distinguish whether there are any orientation specific differences between models, we have also plotted the error and correlation matrices for current flowing along the right-left axis (Figures 3B,E,H) and the inferior-superior axis (Figures 3C,F,I). Sources in the AP direction were omitted here as the field magnitudes are considerably (∼20 times) smaller, but the plots are available in the Supplementary Information. For right-left current flow, we see the relative error (Fig 3B) split into two clusters, the models containing bone and bone-free models. The correlations show high similarity for all realistically shaped models (Fig 3E), before falling for the large sphere and infinite models. With the two metrics overlaid (Fig 3H), we see the bone-free models (relative to the FEM) have large errors, but curiously high correlation - implying the field patterns are similar but overall gain has been affected. Sources oriented inferior-superior show overall low errors between models and high correlations (especially for any models realistically shaped such as the corrected sphere up to the FEM).

### 3.2 Field decomposition reveals orientation-specific sensitivity for bone models

For each source in the medial location along the spinal cord, we performed SVD on each triplet of lead fields for a given location to determine which principal orientations of current flow each volume conductor model is sensitive to. Figure 4A and 4B show examples of the first two eigenfields for a model that includes no bone (1c LC; Fig. 4A) and for a model with bone (5c LC. Fig 4B). These two models were chosen as they employ the same numerical approach to solve the forward problem (linear collocation BEM). The eigenfield topographies plotted have been un-normalised (i.e. we are plotting the columns of ***US***) to visualise their relative intensities. In the bone-free 1c LC model (Fig. 4A), the first component represents current flow oriented along the right-left axis. In other words, when all directions of current flow are equally represented in the cord, the predominant picture at the sensor level is that of current flowing left-right. The second component, by contrast, would correspond to current flow in the superior-inferior direction. The two components have similar intensities (the ratio between the two eigenvalues contained in ***S*** is 1.07). The third eigenfield (not pictured) represents a current flow in the anterior-posterior direction and is (as expected) considerably smaller than the other two components (eigenvalue 3 is 27 times smaller than eigenvalue 1). If we include a bone component in our volume conductor modelling (Fig. 4B) we observe two differences, first the preferential order of the orientations is now superior-inferior followed by left-right. Second the intensity of the second component is considerably smaller (ratio of eigenvalue 1 and eigenvalue 2 is 6.49).

**Figure 4.**
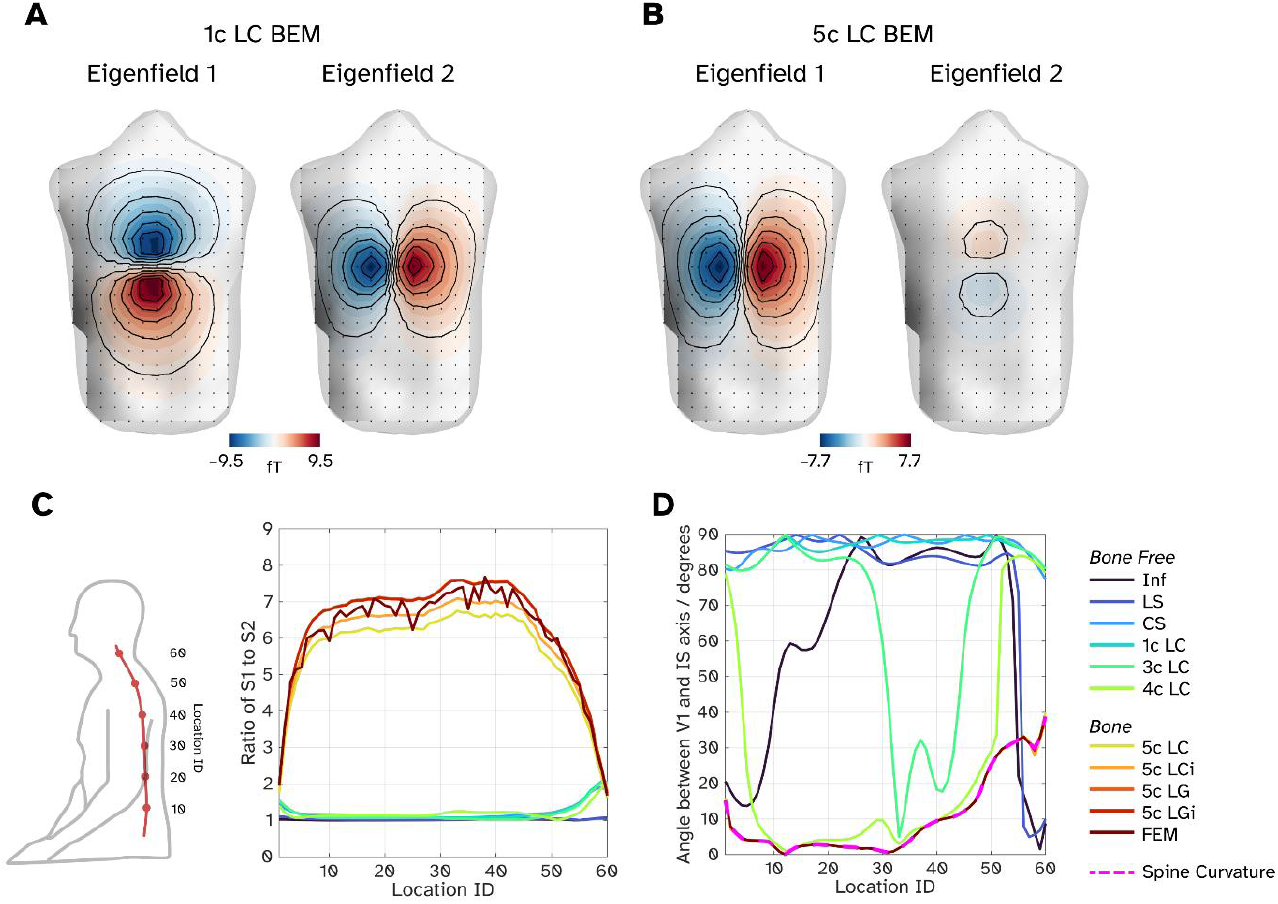
Analysis of sensitivity to the orientation of current flow but modelling source currents in all three cardinal axes and decomposing the field topographies **A)** First two eigenfields for a source located approximately at T9 for a 1-comparment model solved with a linear collocation BEM, showing similar sensitivity in the first two primary axes. **B)** First two eigenfields for the same source but using a 5-compartment model solved with a linear collocation BEM, revealing one dominant component along the inferior-superior axis. **C)** Quantification of the relative strengths of the first two eigenfields for a given model and source by measuring the ration of the two associated eigenvalues. Lower location ID indexes the inferior-superior (lower-higher values) position along the cord (see left plot of panel for a guide). **D)** The angle between the inferior-superior axis and the first eigenfield, the magenta dashed line represents the angle between the spine and the inferior-superior axis.

We quantify these differences in orientation sensitivity for all models across all medial sources in Figures 4C-D. Here, the source IDs are such that lower IDs represent the inferior/lumbar portion of the spinal cord and superior portions have higher IDs. Figure 4C shows the ratio between the first and second eigenvalues at each medial point of the spinal cord. Between sources 10-50 (where we think the whole field topography is adequately sampled) the bone-free models (cooler coloured lines) have a ratio of approximately 1, whereas the models with bone (warmer colours) have a much higher ratio (min: 5.92, max: 7.70). Figure 4D shows the angle between the inferior-superior axis and the orientation of the source that would produce the dominant eigenfield. For example, in Figure 4A, the dominant current flow is left-right and therefore, there is a 90 degree angle to the anterior-superior direction; in 4B by contrast the angle is zero. Figure 4D shows that for bone free models, the dominant field pattern is often due to the source component in the left-right direction. Adding additional compartments (3c-4c LC) shows that these tissues start to dictate the primary sensitivity axis (as the first two eigenmodes have similar amplitudes). For the 3c LC model, we see that the lungs and heart affect sources with IDs 30-50, with the primary orientation rotating to be nearer the IS axis rather than the RL axis. Adding the 4^th^ compartment (white matter of the spinal cord) shows the primary sensitivity axis is closer to the curvature of the spine (dashed magenta line). However, in both these cases, the first and second eigenfields are of similar intensity and so this preferential orientation discussion is a moot point. As soon as we add the bone into the models, we see that the primary eigenfield is oriented along the direction of the spinal curvature. Combined with the fact the first eigenfield is several factors larger than the second, the bone appears to give maximal sensitivity along the **cord’s** superior-inferior axis, at the expense of all other orientations.

### 3.3. Bone-based models may offer better lateral source separation

Finally, we compare the field topographies of the left medial to the right medial sources for a given slice of spinal cord. The motivation here is to determine if the field topographies offer enough separability between the two sources to determine the lateralisation of a current, so we are looking for higher relative errors and lower correlations between a pair of topographies. Here, we specifically compare current flow oriented along the curvature of the spine to reduce our comparisons down from 3 field topographies per source to 1. Figure 5A shows boxplots representing the relative errors between pairs of field topographies, with each jittered point representing a pair of sources. Increasing the complexity of bone-free models (Inf to 4c LC) lead to larger relative errors between the left and right sources, but it is the introduction of the bone compartment which leads the largest increase of errors between sources. Given the field topographies are of a pair of sources should be approximately equal in magnitude, we would expect the correlations between fields to be inversely proportional to the errors. This is what we observe in Figure 5B, where bone-based models tend to have lower squared-correlations between sources. To illustrate one case, Figures 5C and 5D show the modelled field topography for a pair of sources in the T9 location and the difference between patterns for a bone-free (1c LC; Fig 5C) and bone (5c LC; Fig 5D) models. The 1c LC model generates field topographies where the extreme fields at the poles are almost equal in magnitude (9.6 fT/nAm v -9.2 fT/nAm, 95% for the left medial source) whereas the bone introduces an asymmetry (7.0 fT/nAm v -8.3 fT/nAm, 84% for the left medial source). This asymmetry in the field patterns and intensities lead to greater differences (or improved discriminability) between the two sources and thus larger relative errors and lower correlations.

**Figure 5.**
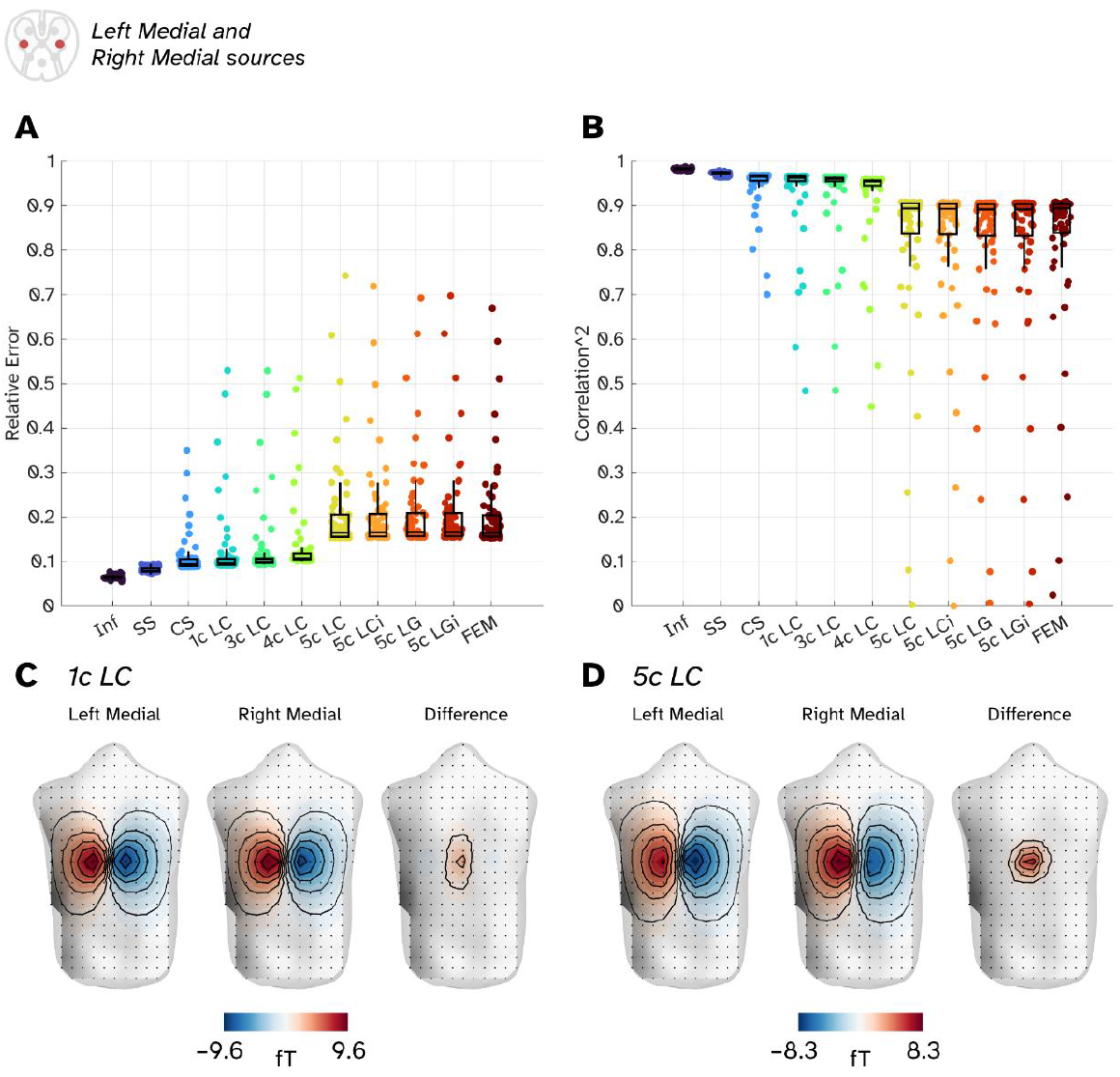
The similarity and differences of field topographies from lateralised sources in the spinal cord. A) Relative errors between pairs of sources in a given slice of spinal cord (left medial and right medial sources). Individual points represent a pair of sources whilst boxplots show the median errors across the entire spinal cord. B) Squared correlation coefficients between pairs of sources. C) Visualisation of the field topographies for a pair of 1 nAm source currents in the T9 region of the spine and the difference between them for a bone free model (1c LC BEM). D) Field topographies for two lateral sources in T9 for a volume conductor containing bone (5c LC BEM).

## 4) Discussion

We have tested different forward models describing how current flow in the spinal cord manifests as magnetic fields outside the torso. We tested these for elements of current flow situated from cervical to lumbar regions of the spinal cord oriented in three directions.

We found that most volume conductor models can approximate the field topography for a source current oriented along the inferior-superior axis of the body (assuming some attempt to model the torso boundary has been implemented). The infinite homogenous medium model generates lead fields which are the most distinct from any other volume conductor tested but, perhaps surprisingly, a large sphere compared well to realistically shaped torsos. Our assumption is that by moving the origin so far forward, all dipoles oriented along the inferior-superior and left-right axes are considered tangential sources (relative to the sphere) origin and so are preserved. In fact, if one continued to move the origin an infinite distance in front of the subject one would converge on a half-space model^26^, where the model approximates into a flat plane representing the subject s back. The corrected sphere was the most similar of the three analytical models to the numerical methods (BEMs and FEM). However, we note that the implementation of the corrected sphere here uses a basis set which does not tolerate the sensors being so close to the conductivity boundary^27^, and so produces artefactual field patterns (see Supplementary Information). So, if one wanted the most rudimentary volume conductor to perform a basic source analysis along the inferior-superior axis, a single compartment boundary element model could be the safest recommendation.

We have used established solvers used in EEG / MEG (and ECG, MCG) and existing geometries to probe how we could approach the forward modelling in magnetospinography. We have chosen to use approaches and techniques commonly used in this field, preferably implemented in EEG/MEG analysis toolboxes. The piecewise homogeneous models that contain only main structures should be rather straightforward to implement for individual geometries, and our template model can also be warped to individual geometry. This work motivates more detailed and more realistic volumetric models of anatomy; we are not aware of any academic software pipelines currently available.

Along the inferior-superior axis, we determine that every nAm of a current dipole gives rise to a maximum absolute magnetic field of around 10 femtoTesla along the surface of the back (assuming the source current is 50 mm deep). This accords with empirical averaged evoked response recordings from SQUID based systems in which sources of around 4 nAm give rise to field changes of approximately 40-50 fT^15^. Current SQUID and OPM-based systems have a white noise floor of around 5 and 20 fT per square root Hertz respectively. This implies that in order to achieve 0 dB SNR evoked response in 100Hz bandwidth, a single trial of SQUID or sixteen trials of OPM recording would be required.

Our headline finding was that the inclusion of bone wrapped around our spinal cord affects the field topographies in non-trivial ways. First, we found that the sensitivity to different orientations of current was driven primarily by whether bone was included in the volume conductor. Bone-free models had equal sensitivity to sources oriented along the superior-inferior and left-right axes, while the bone-inclusive models had a single preferential axis: along the curvature of the spinal cord. The second component (left-right oriented) is attenuated in magnitude in our simulations by as much as 5- to 8-fold, depending on how well sampled the source is by the sensors and which forward solution was implemented. The field topographies of left-right oriented sources produced were similar in pattern across all numerical models (BEMs and FEM), implying that in our simple model of the vertebrae, the bone essentially acts as a “global” attenuator of the sensitivity to current flow off-axis to the inferior-superior axis of the spine. When separating out the effects of primary and secondary current flow on the magnetic field, we see that the secondary currents generates an almost equal and opposing field to the primary currents for transverse oriented current flow (see Supplementary Information). The proportion of attenuation is (in part) dependent on the selection of conductivity values employed to represent the spinal cord and bone meshes. The larger the ratio of conductivity between the two interfaces, the larger the attenuation of off-axis sources (see the 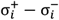 term in Equation 3, and the simulations in the Supplementary Information). We note that our bone model was not anatomically accurate (vertebrae are not rotationally symmetric, they are discrete entities, and they vary in size) and, in practice, this effect may be even more complicated to characterise than merely a simple scaling of the field strength. we would still expect there to be an orientation dependent effect with a realistic geometry. This work is based on a single idealized participant. Future work might look at the intrinsic variability expected in forward models across participants and how much additional complexity is required.

Our second bone-based model observation is how the inclusion of bone allows for theoretically better separation of sources in the same transverse plane of the spinal cord. We found that sources placed 8 mm apart in the same transverse slice of spinal cord had associate field topographies which were more dissimilar if bone was incorporated. In part, this was due to how poles of the field topography were distorted by bone. The pole proximal to the bone walls attenuated more than the other. We note that the similarity in field patterns based on correlations was still very high (around 90 % variance explained by each other) in our theoretical sensor array, but with methods available to optimise sensor sampling^40 42^ we may reduce the correlations further. The use of bone-based models may allow for direct confirmation of source laterality in the spinal cord, something only fMRI has demonstrated (non-invasively) so far^43,44^.

Ultimately, these simulations reveal that we need *a priori* knowledge of the location of the spinal cord and spinal column that surrounds it for precision imaging of spinal cord sources. This is especially critical for current flow which does not flow along the cord in inferior-superior orientation (from for example, integration from spinal interneurons^45^). Given that current flow in the cord along the inferior-superior axis is much simpler to model one could derive the depth of the spinal cord from observed sensor-level data using generative modelling approaches (something which has been investigated with electrophysiological recordings from the brain^46^). For off-axis current flow however this quickly becomes computationally expensive if we must solve a complex BEM or FEM volume conductor every time we need to iterate our source space, as we would also need to iterate the morphology of the spinal cord and bone compartments. SQUID-based MSG systems have taken an X-ray of the participant in situ of the sensors^17,47,48^ to determine their source space. From a practical perspective, future work might consider leveraging anatomical model derived from the individual spine and torso with (for example) MRI^49^. However, from a practical perspective this is by no means a trivial thing to do. Unless carefully controlled, the participant’s posture (hence the position of spinal cord), will change between scanning modalities, and potentially undermine any gains in complexity-something that increasingly accurate forward models cannot tolerate well^20,50^. We made use of a generic sensor array based on a specific type of OPM sensor at fixed spacing. Future work might look at the relative advantages of sensor arrays which could become more granular (to better sample superficial sources) or closer to the torso (to increase sensitivity).

In summary the inclusion of the bone in volume conductors for source currents in the spinal cord generates features which cannot trivially be emulated in the absence of bone, and so the inclusion of the bone will be essential to maximise the quality of the source analysis MSG can provide. With this in mind, we can now focus on both better methods to image and constrain the anatomy of the spine to generate plausible volume conductor models and optimise sensor placement for denoising recordings and inverse modelling in the future.

## Acknowledgments

GCO is funded through an UKRI Frontier Research Grant [EP/X023060/1] and acknowledges EPSRC [EP/T001046/1] funding from the Quantum Technology hub in sensing and timing (sub-award QTPRF02). MES is supported by a Wellcome Technology Development grant [223736/Z/21/Z]. MS is supported by the UZH Global Strategy and Partnerships Fund Scheme. SM was supported by an EPSRC Healthcare Impact Partnership Grant [EP/V047264/1] and acknowledges support from the Federal Commission for Scholarships for Foreign Students for the Swiss Government Excellence Scholarship (ESKAS No. 2024.0251). This research was supported by the Discovery Research Platform for Naturalistic Neuroimaging funded by Wellcome [226793/Z/22/Z].

## Data availability

Code for generation of the results can be found at https://github.com/georgeoneill/study-spinevol, with an archived version of the code (and all dependencies) available at https://doi.org/10.5281/zenodo.14883493. Supporting toolbox for the generation of the volume conductors and solvers can be found at https://github.com/fil-opmeg/torso_tools. The linear Galerkin BEM solver is property of Matti Stenroos (Aalto University) and is not publicly available. For collaborations involving the Galerkin solver, please contact Matti Stenroos.

## Notes

### Competing Interest Statement

The authors have declared no competing interest.

### Summary of Updates

This version has been revised according the reviewer's requests. Major changes include a more detailed theory section, and updated results based on improved meshing for finite element analysis.

