## Supplementary Information for "Volume conductor models for magnetospinography"

George C. O'Neill[a✉], Meaghan E. Spedden[b], Maike Schmidt[b], Stephanie Mellor[b,c,d], Matti Stenroos[e], Gareth R. Barnes[b✉]

[a] Department for Neuroscience, Physiology and Pharmacology, University College London, London, UK
[b] Department of Imaging Neuroscience, UCL Queen Square Institute of Neurology, University College London, London, UK
[c] Spinal Cord Injury Center, Balgrist University Hospital, Zurich, Switzerland
[d] Translational Neuromodeling Unit (TNU), Institute for Biomedical Engineering, University of Zurich & ETH Zurich, Zurich, Switzerland
[e] Department of Neuroscience and Biomedical Engineering, Aalto University School of Science, Espoo, Finland
✉GCO: g.o'; GRB:

---

Contents:

1) Field comparison results (continued)
2) The effect of adjusting the conductivity of the bone
3) Visualisation of the magnetic field due to volume currents
4) The effect of sensor proximity on the corrected sphere model
5) Assessing the mesh density for convergence of BEM solutions
6) Field plots from the same source for all models

1) Field Comparison Results (continued)

In the main manuscript we compared the errors and similarities between all forward models, but excluded an analysis on sources which were oriented along the anterior-posterior axis as these sources explain only a tiny proportion of the variance in the generated field topographies. For completion, we have included the results for the medial sources oriented along this direction, these can be found in Figures S1A-C. We see that the three clusters of bone-based, bone free numerical and bone free analytical conductive models persists.

Given the finding that the bone-based models had the most sensitivity along the curvature of the spine, we also repeated these measures for dipoles oriented which ran parallel to the curvature of the spine itself. The results of this are in Figure S1D-F.

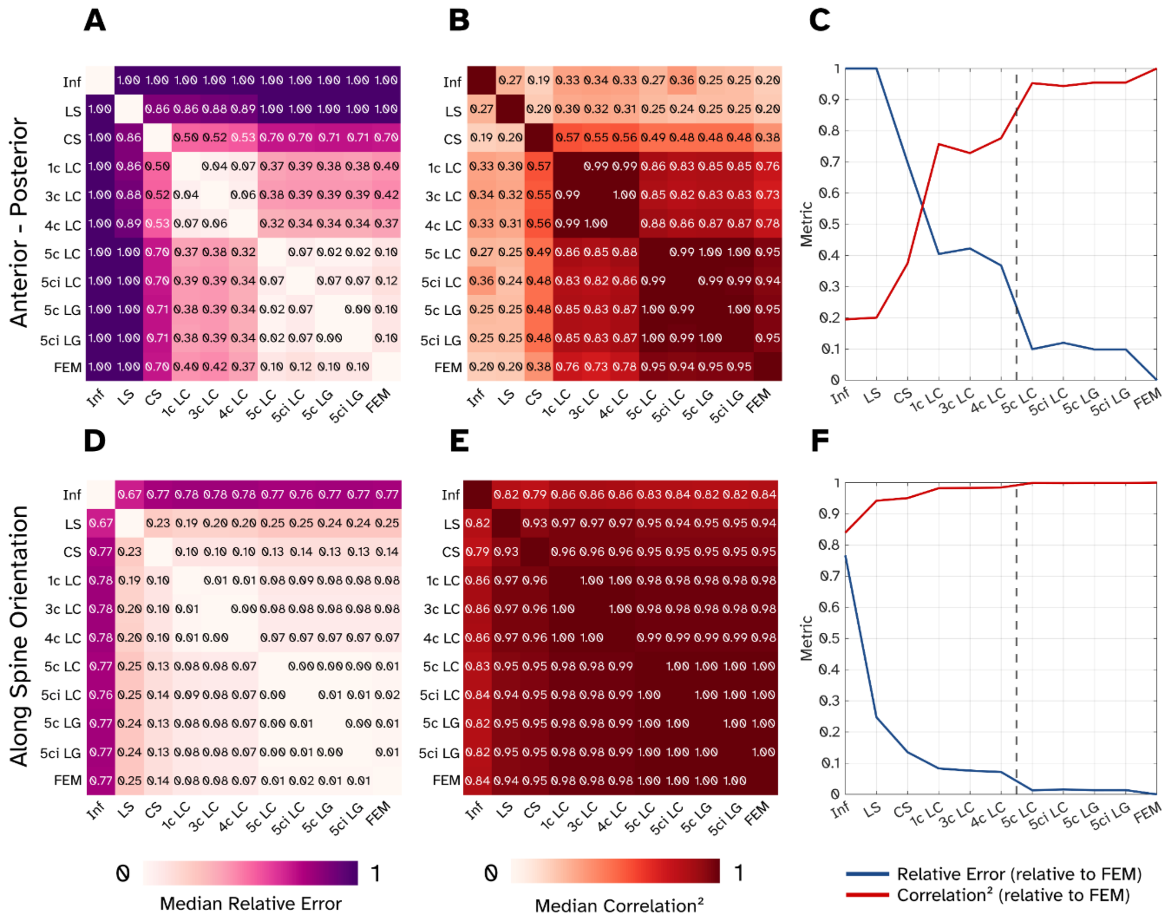


Figure S1 – Field comparison results between models for sources oriented along anterior-posterior directions (panels A-C) and parallel along the orientation of the spine's curvature (panels D-F).

### 2) The effect of adjusting the conductivity of the bone

In the main manuscript, Equation 3 shows that the volume field produced is proportional to the change in conductivity across a boundary interface (e.g. from the spinal cord to the bone). We demonstrate this numerically here by adjusting the conductivity of the bone mesh relative to the spinal cord mesh (which as a reminder is set to 0.33 Sm-1). We set the bone conductivity to be 0.33/n, where n was set to be a range of values between 1 and 40 (40 was the value used in the main manuscript), Figure S2A compares all the lead fields generated relative to when n (which we call the conductivity ratio in the figures) to 40. As the conductivity ratio between the two boundaries falls (i.e. the bone becomes more conductive) the errors relative to a ratio of 40 increase, but the correlations remain high. Indicating this is generating (mostly) magnitude related changes. Figure S2B we performed the eigenfield analysis similar to the main manuscript to understand the ratio of variance between the first two components for a given source. Here we see as the conductivity increases, the ratio for the first two eigenvalues increases too. Note this this is almost a linear effect.

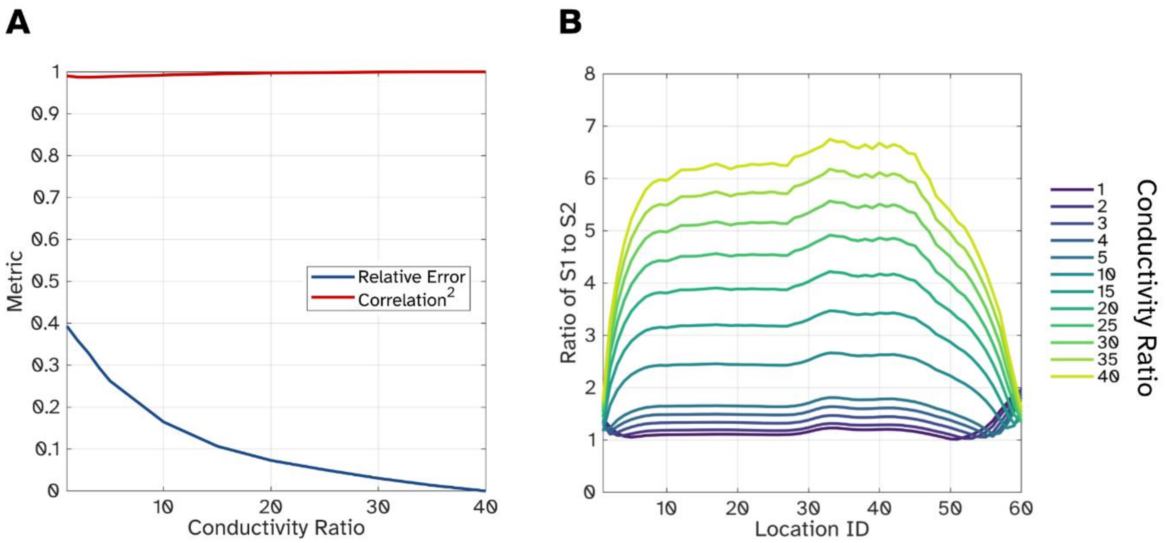


Figure S2 – The effect of adjusting the conductivity of the bone in the 5c LC model relative to the spinal cord mesh, here we report the ratio of connectivity between the spinal cord and bone. A) The correlation and relative errors of all modelled sources relative to the model used in the paper. B) The ratio of the first two eigenvalues from the eigenfield analysis.

### 3) Visualisation of the magnetic field due to volume currents

Because we are operating in the quasistatic regime (i.e. any oscillatory fields are slow enough to ignore the time dependence of Maxwell's equations), the combination of the magnetic fields arising from the primary current flow and returning volume currents becomes a simple sum[1],

$$\boldsymbol{B} = \boldsymbol{B}_{prim} + \boldsymbol{B}_{vol}. \qquad (S1)$$

In Figure S3 we have resolved the field components into the primary and volume fields for two volume conductors (1c and 5c LC BEMs) and two dipole orientations (right to left, inferior to superior). We see that for the inferior to superior oriented current flow, the volume fields for both the 1c and 5c BEMs are similar in magnitude and topography, whereas for the right-to-left current flow these are quite distinct from each other. The 5c BEM volume currents produce a field which is almost identical to the primary field but in the opposite direction (correlation coefficient: -0.88).

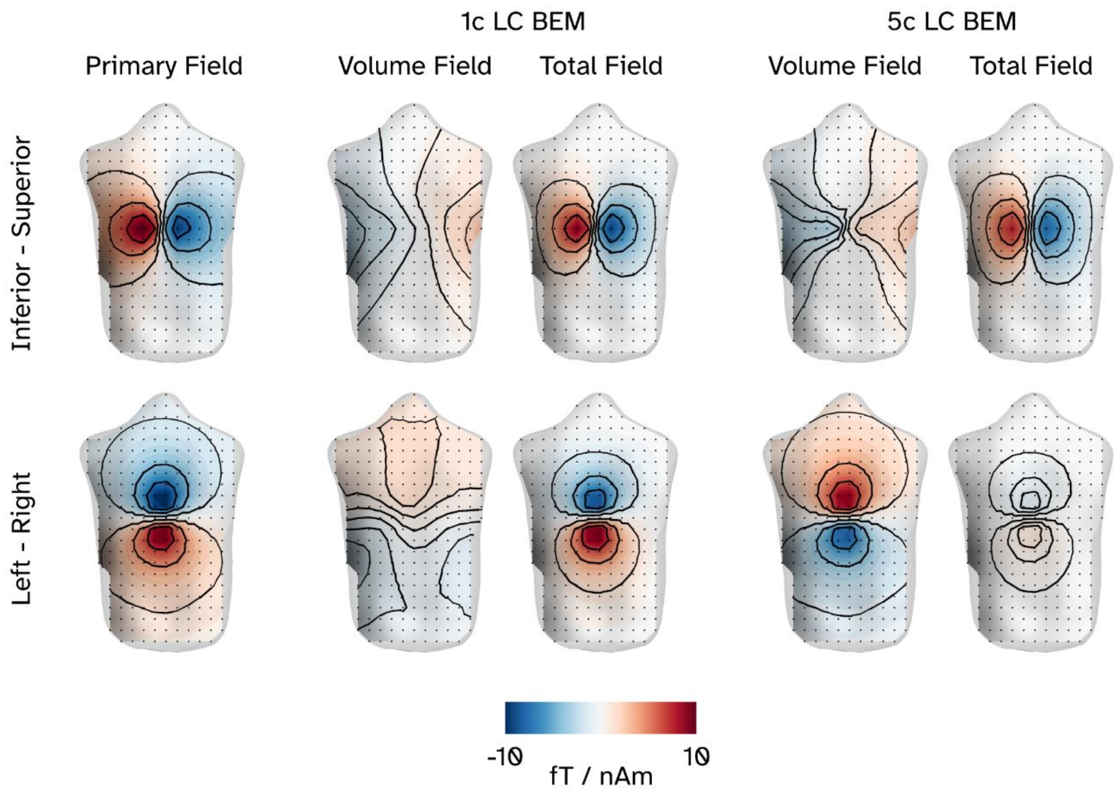


Figure S3 – The separation of the magnetic field components due to the primary and volume currents.

4) The effect of sensor proximity on the corrected sphere model

In the original paper describing the corrected sphere model[2], there is discussion on how the current basis set is ill-suited to model the local, focal field components near the boundary. We demonstrate below that this has tangible effect on the field topographies produced. We simulated sources along the spine with the same theoretical array from the main manuscript, but this time we shifted the distance between sensors and boundary to range from a (physically implausible) 1 mm up to 40 mm away. We compared the field topographies from the Infinite, Large Sphere, Corrected Sphere and 1c and 5c BEM models. Figure S4A shows how the correlations between the 5c BEM and the other models is affected by the distances between the torso boundary and sensors. Over the range of distances the Infinite, Large Sphere and 1c BEM models have a similar relationship to the 5c BEM. However we see for the Corrected Sphere (Fig. S4A yellow line) there is a monotonic decrease in similarity to the 5c BEM the closer the sensors get. For the main study we fitted spherical harmonics up to an order of $\ell$=10, which may be an acting as a smoothing kernel on the true higher frequency components. To determine if increasing the order would improve the accuracy, we adjusted the basis set to fit from $\ell$=0-25 (i.e from a sphere up to a more complex representation). We see in Figure S4B, for a “sensible” distance (such as 10 mm or further) this can offer modest improvements in the accuracy, but for closer distances this does not. For comparison, we plot the field topographies for the Corrected Sphere and 1c BEM for a range of distances in Figure S4C (note the intensity of the field topographies are arbitrary here). The BEM field patterns, even at close proximity to the sensors produce smooth field topographies and predictable isolines compared to the Corrected Sphere.

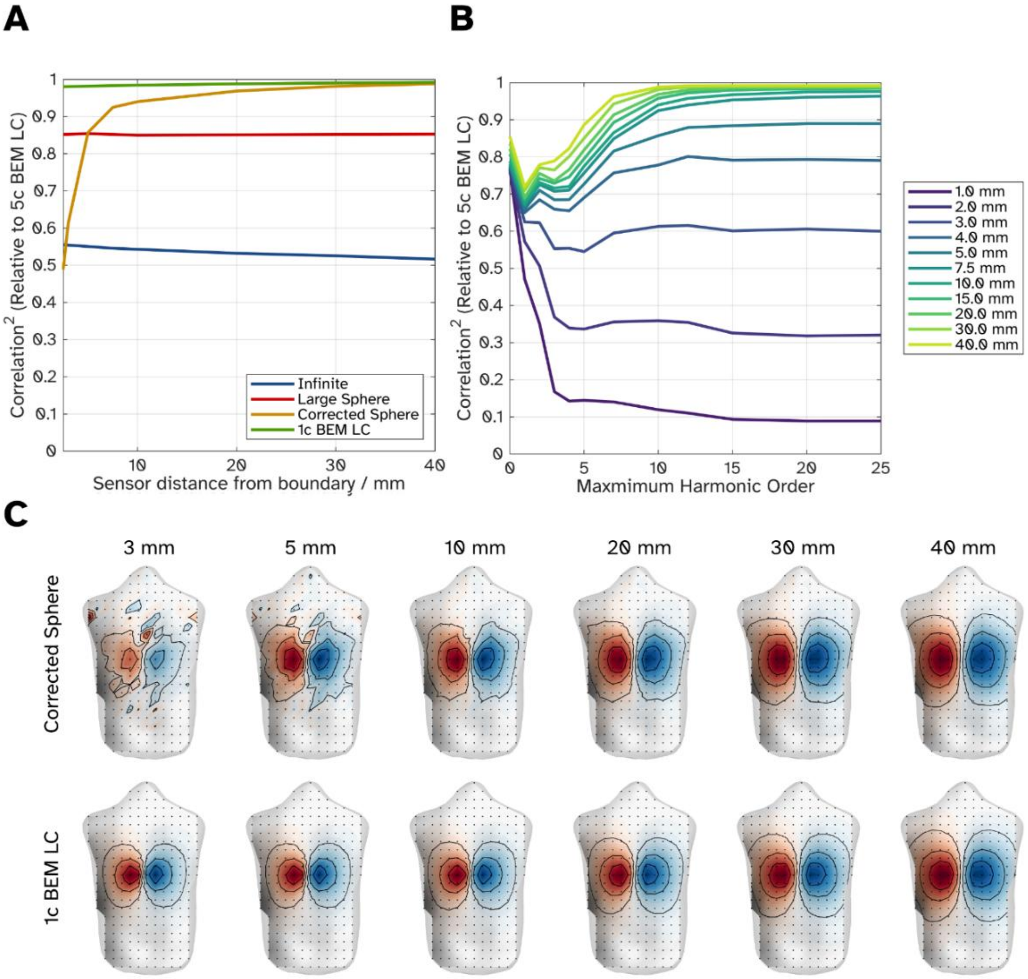


Figure S4 – The effect of the distance between the sensors and torso boundary on different models. A) The correlation between the 4 simplest models and the 5c BEM as a function of distance. B) The correlation between the Corrected Sphere and the 5c BEM for a variety of distances as a function of the maximum harmonic order modelled. C) A comparison of modelled field topographies for a current source oriented inferior-superior at approximately the T9 point of the spine.

5) Assessing the mesh density for convergence of BEM solutions

To test whether we had sufficiently sampled the meshes used at the boundary conditions, we both oversampled and downsampled the boundary meshes in question to see how much our field topographies altered compared to our original mesh density. Here density refers to the number of faces in the mesh with 100% representing the number of faces in the meshes used in the main manuscript. 400% sampling was achieved dividing each face into 4 smaller faces. Then, we iteratively downsample the number of faces to a target percentage (20-380%) and compare the field topographies generated to the oversampled (400 % density) field topographies.

First, we investigated the simple 1C LC BEM, which consists of a single torso-mesh as the boundary in Figure S5A. We modelled all sources in the outer ring of every "axial slice" of the spinal cord in all three cardinal orientations, generating 1464 field topographies from 488 locations. Figure S5A shows the median correlation (red line) and relative errors (blue line) between the downsampled results and full density results on a log-axis. Compared to the 400 % density mesh, the 100 % torso mesh used in the main manuscript has a median relative error of 0.015 and median correlation of 0.999.

Figures S5B-D show the results of adjusting the densities of the spinal cord and bone meshes for the 5c LC BEM (whilst keeping the torso mesh at 100 % density for computation convenience). Here we use when both the spinal cord and bone meshes have been oversampled to 400 % density as the reference model. Figure S5B shows the results of adjusting both the spinal cord and bone meshes simultaneously. Comparing the meshes in the paper (100 % density) to the oversampled reference model, the median relative error is 0.052, and the median correlation is 0.997. We can adjust the two meshes independently of each other, keeping one fixed at a 400 % density to understand which mesh has more effect on our findings. Perhaps unsurprisingly this is driven by the spinal cord mesh, which as shown in Figure S5C has near identical results (median relative error: 0.054; median correlation: 0.997 at 100 % density), whereas the bone mesh density effect is less pronounced (Figure S5D; median relative error: 0.003; median correlation > 0.999 at 100 % density).

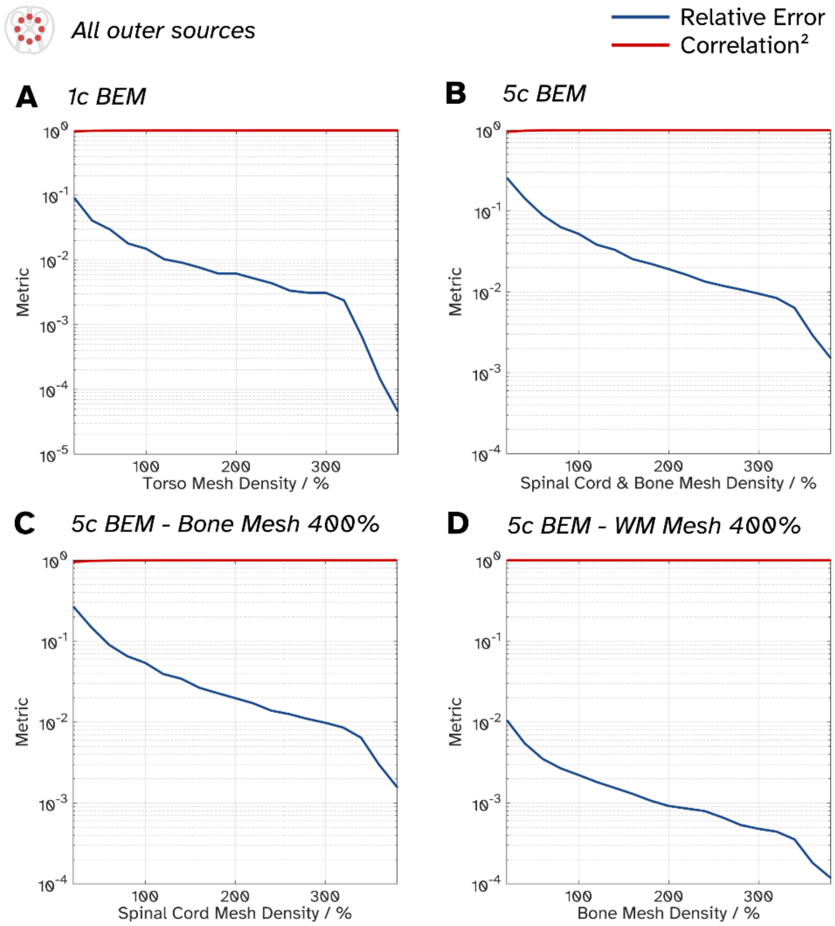


Figure S5 – Comparing the field topographies for boundary elements models between the subdivided triangle boundary meshes (noted as 400 % density), and iteratively down sampled representations. Here 100 % density refers to the meshes used in the main manuscript. A) The median correlation and errors of 1464 field topographies for a 1 compartment boundary element model (1c LC, where the boundary in question was the torso). B) Median errors for the same sources, but instead for the 5c LC model, where the spinal cord and bone meshes are resampled. C) Metrics for 5c LC model, when the bone mesh is held at 400 % density and the spinal cord mesh is adjusted, with the opposite treatment for (D).

6) Field plots from the same source for all models

Figures S6-S16 are the plots for the same source located at approximately the T9 location of the spine for all 11 models tested. Isolines here demarcate the deciles of field strength for a given row.

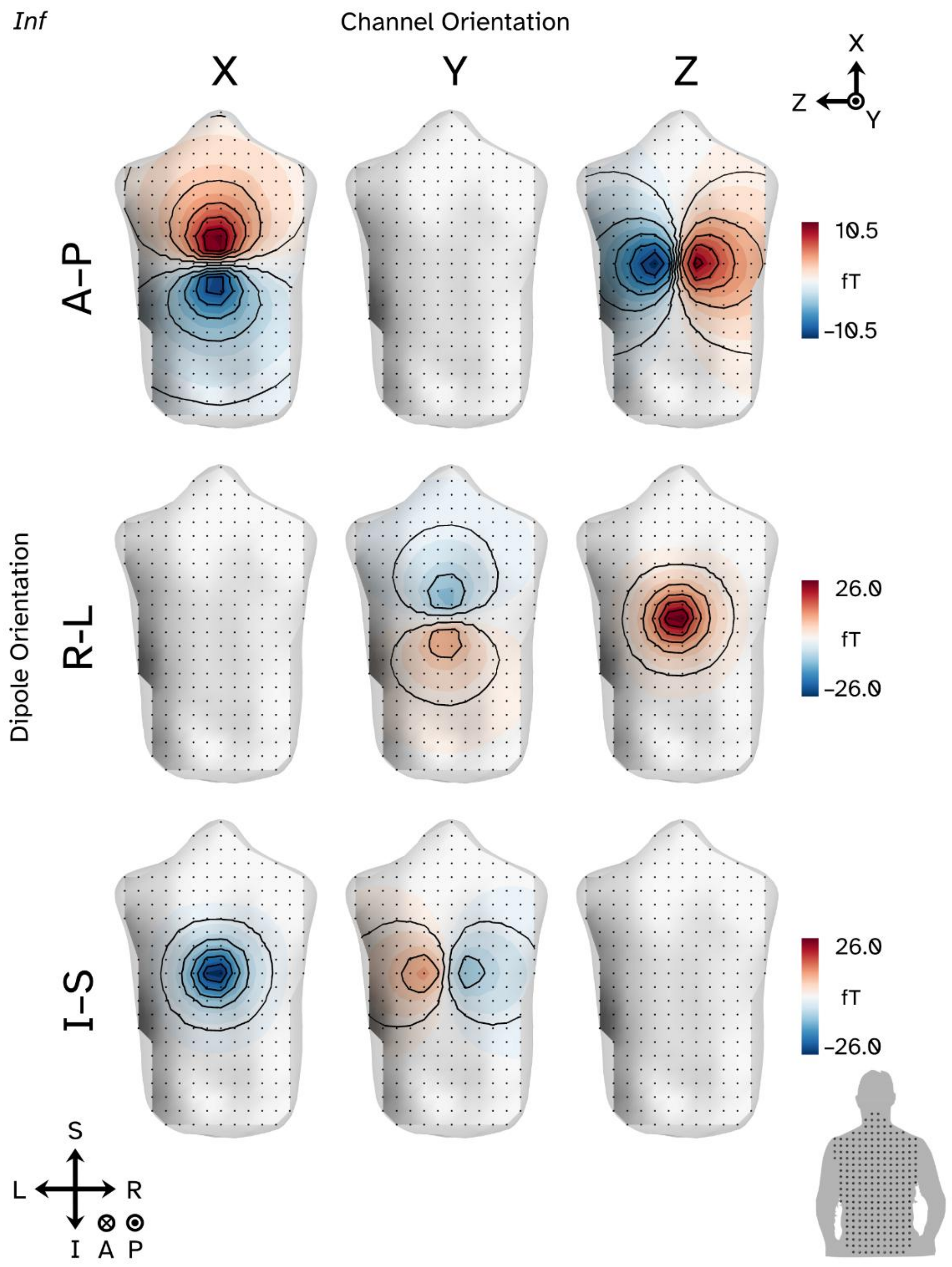


Figure S6

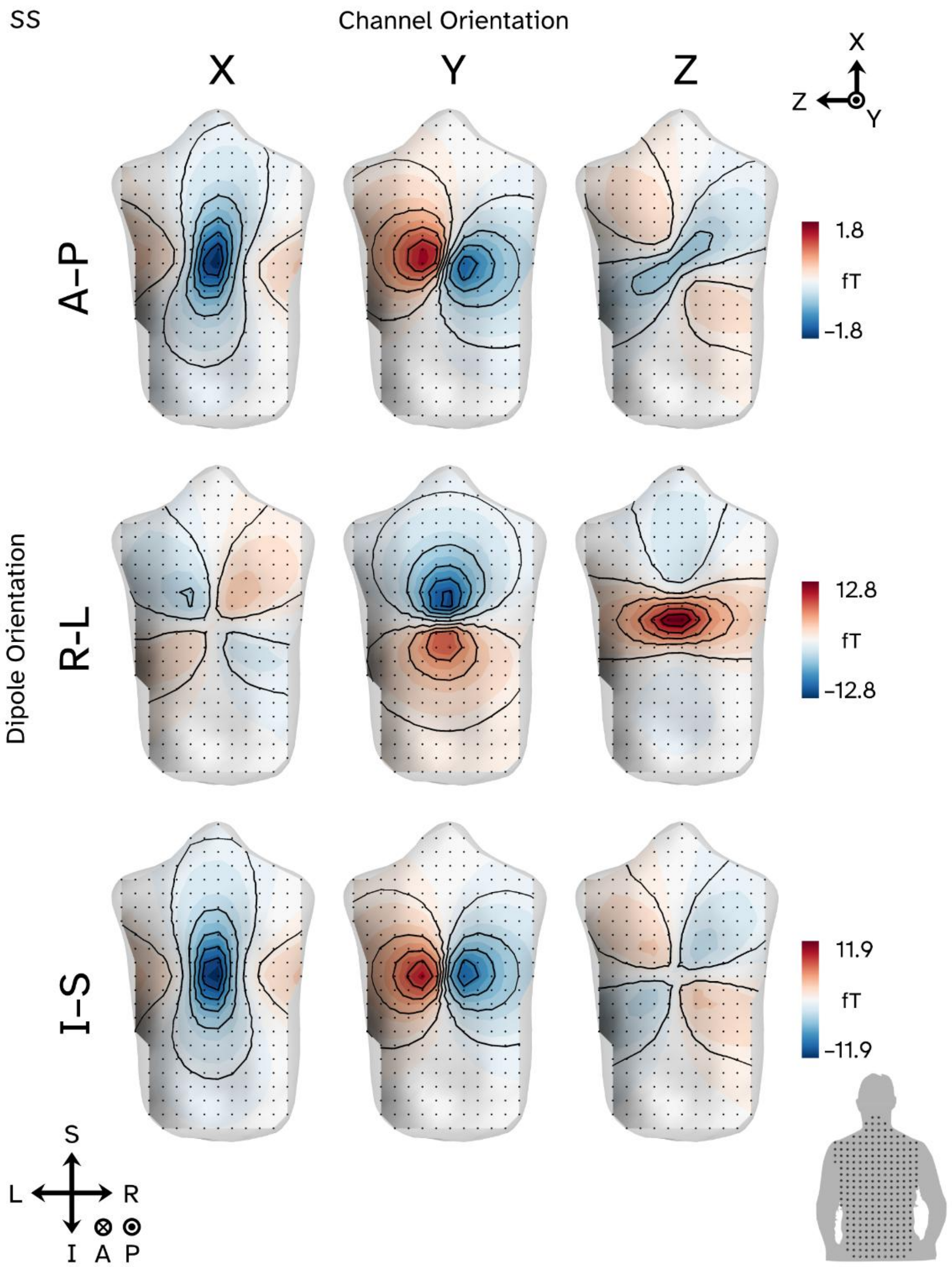
SS
Channel Orientation
X
Y
Z
X
Z
Y
A–P
1.8
fT
−1.8
Dipole Orientation
R-L
12.8
fT
−12.8
I–S
11.9
fT
−11.9
S
L
R
I
A
P

Figure S7

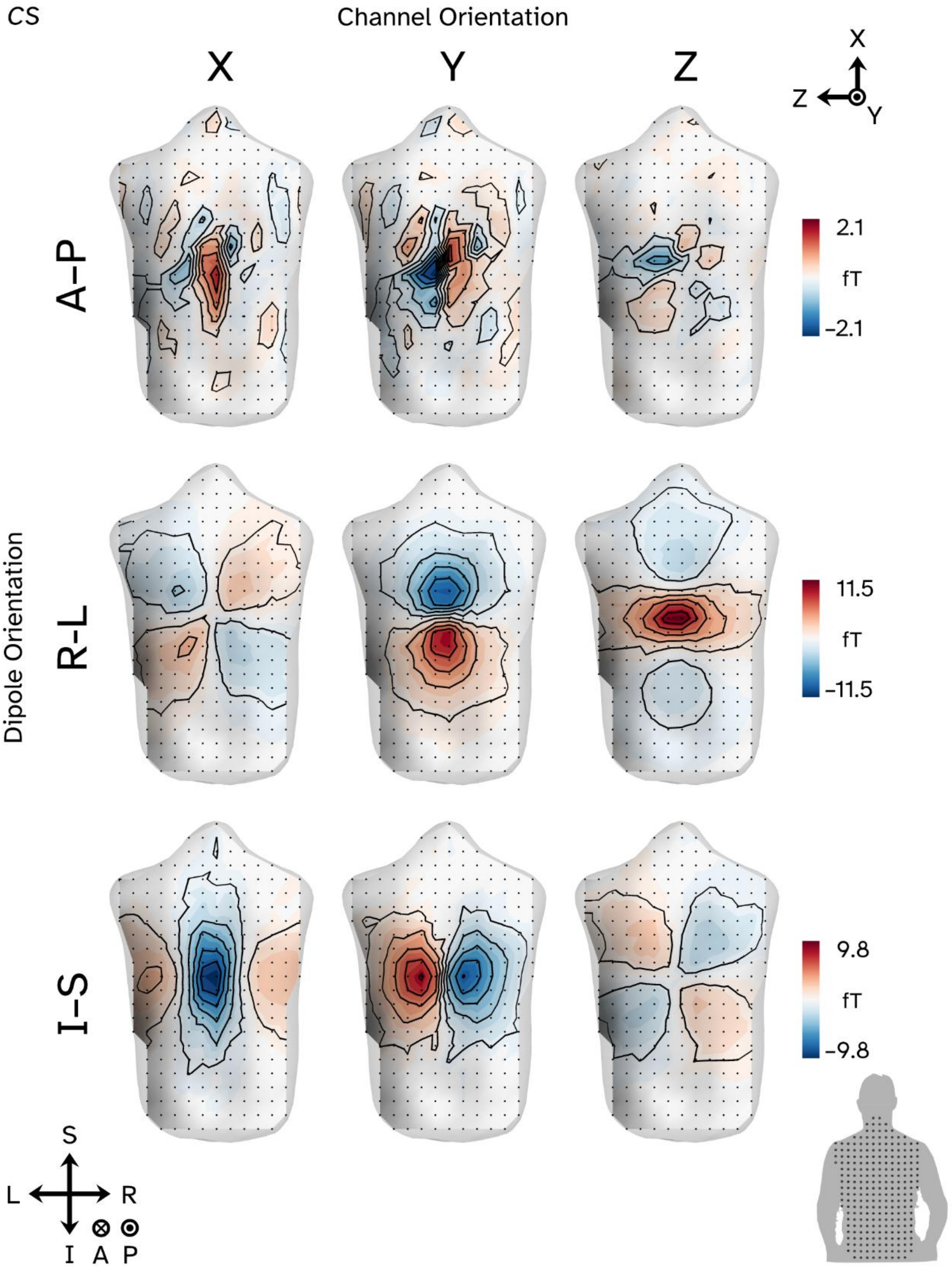
CS
Channel Orientation
X
Y
Z
X
Z
Y
A–P
2.1
fT
−2.1
Dipole Orientation
R–L
11.5
fT
−11.5
I–S
9.8
fT
−9.8
S
L
R
I
A
P

Figure S8

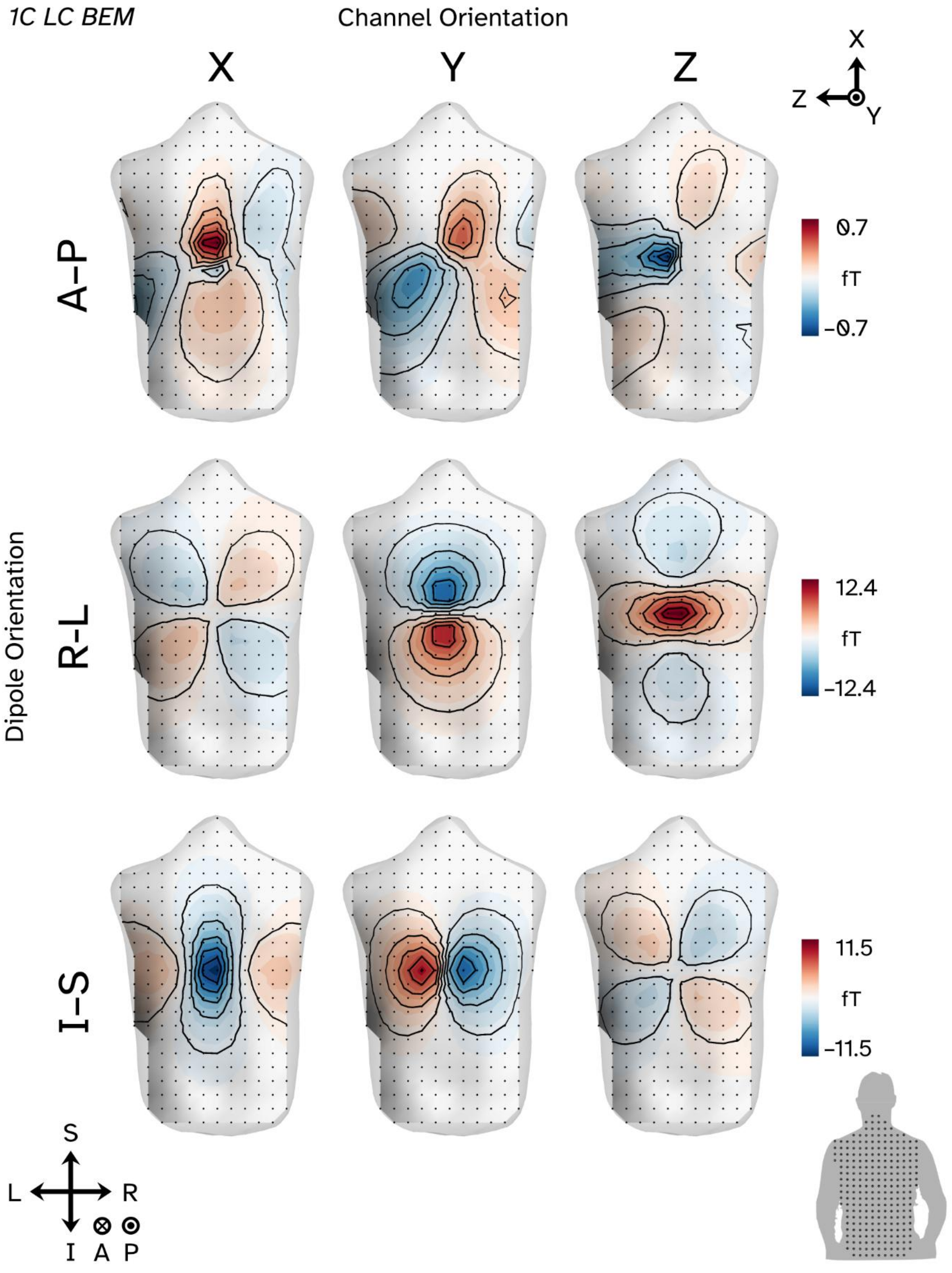
1C LC BEM
Channel Orientation
X
Y
Z
X
Z
Y
Dipole Orientation
A–P
R–L
I–S
0.7
fT
−0.7
12.4
fT
−12.4
11.5
fT
−11.5
S
L
R
I
A
P

Figure S9

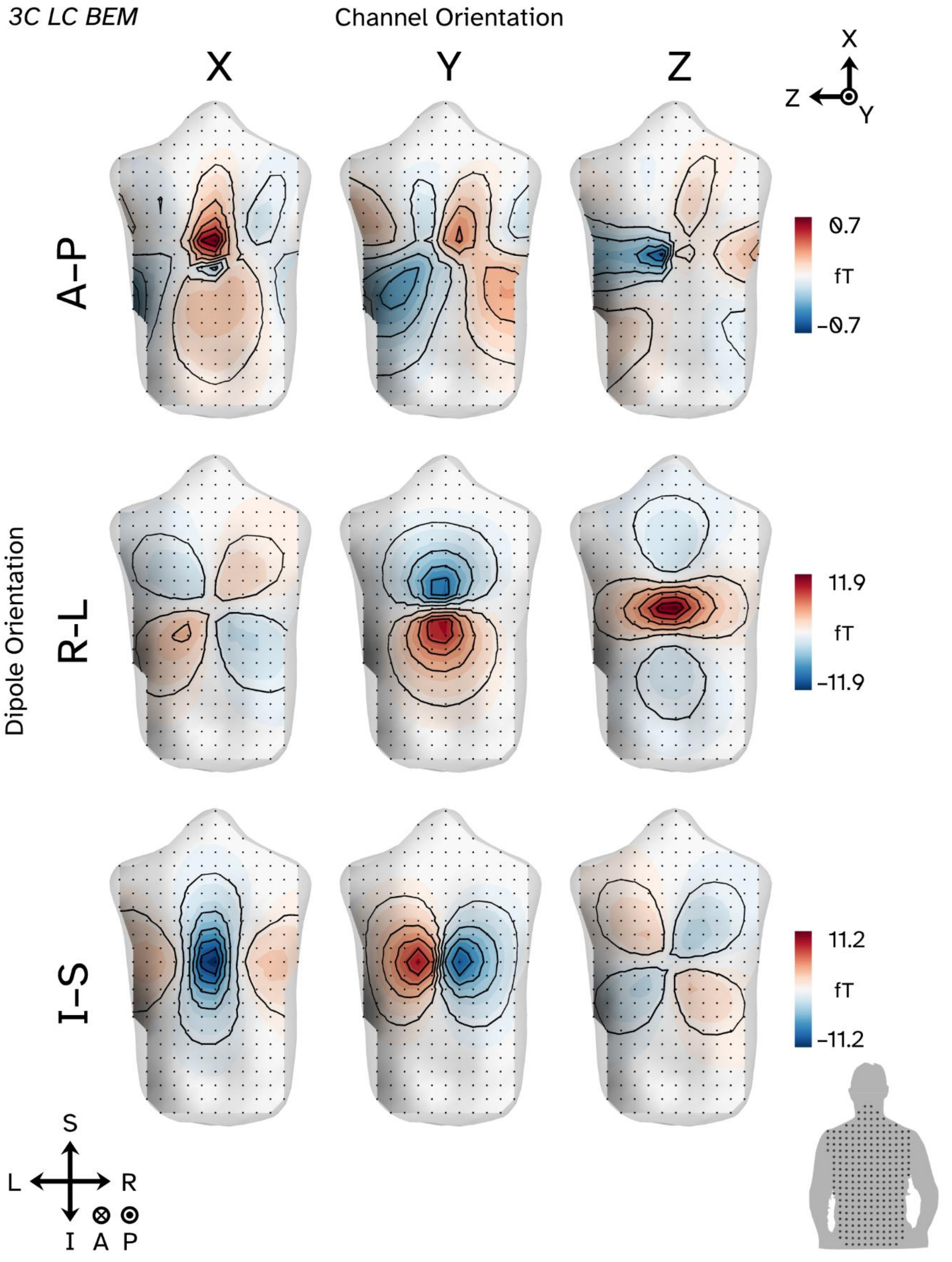

3C LC BEM
Channel Orientation
X
Y
Z
X
Z
Y
Dipole Orientation
A–P
R–L
I–S
0.7
fT
−0.7
11.9
fT
−11.9
11.2
fT
−11.2
S
L
R
I
A
P


Figure S10

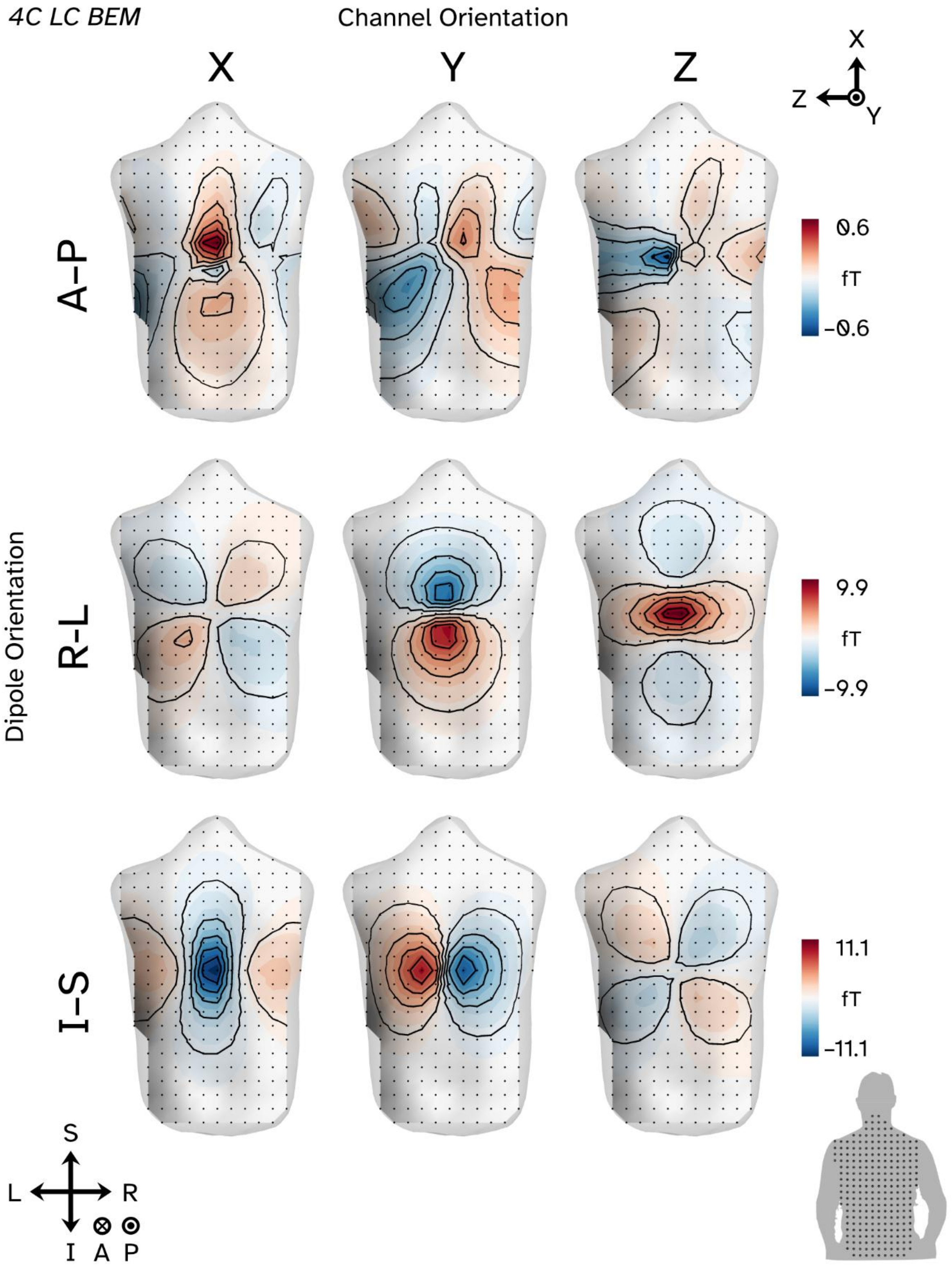
4C LC BEM
Channel Orientation
X
Y
Z
X
Z
Y
Dipole Orientation
A–P
R–L
I–S
0.6
fT
−0.6
9.9
fT
−9.9
11.1
fT
−11.1
S
L
R
I
A
P

Figure S11

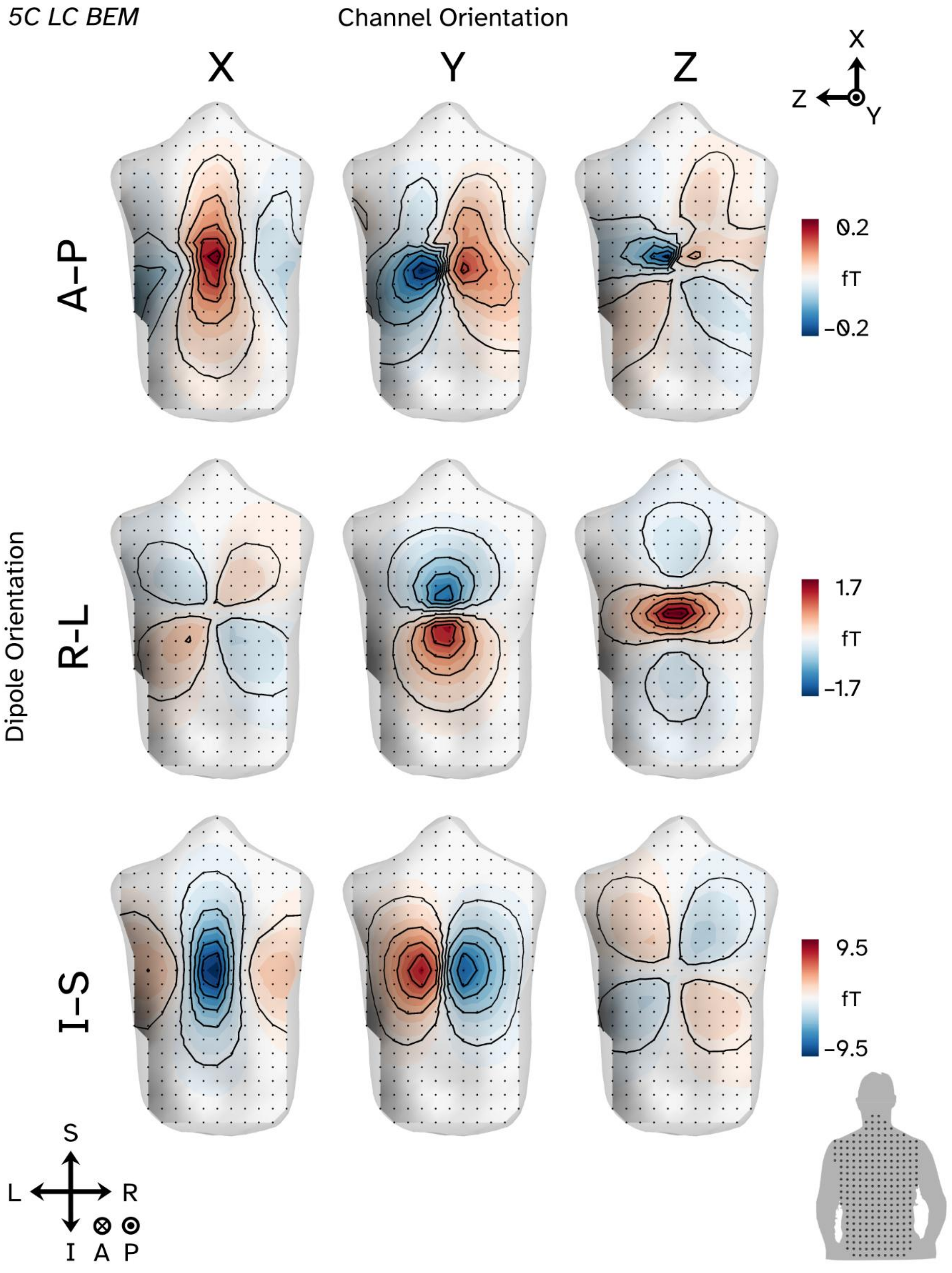
5C LC BEM
Channel Orientation
X
Y
Z
X
Z
Y
Dipole Orientation
A–P
R–L
I–S
0.2
fT
−0.2
1.7
fT
−1.7
9.5
fT
−9.5
S
L
R
I
A
P

Figure S12

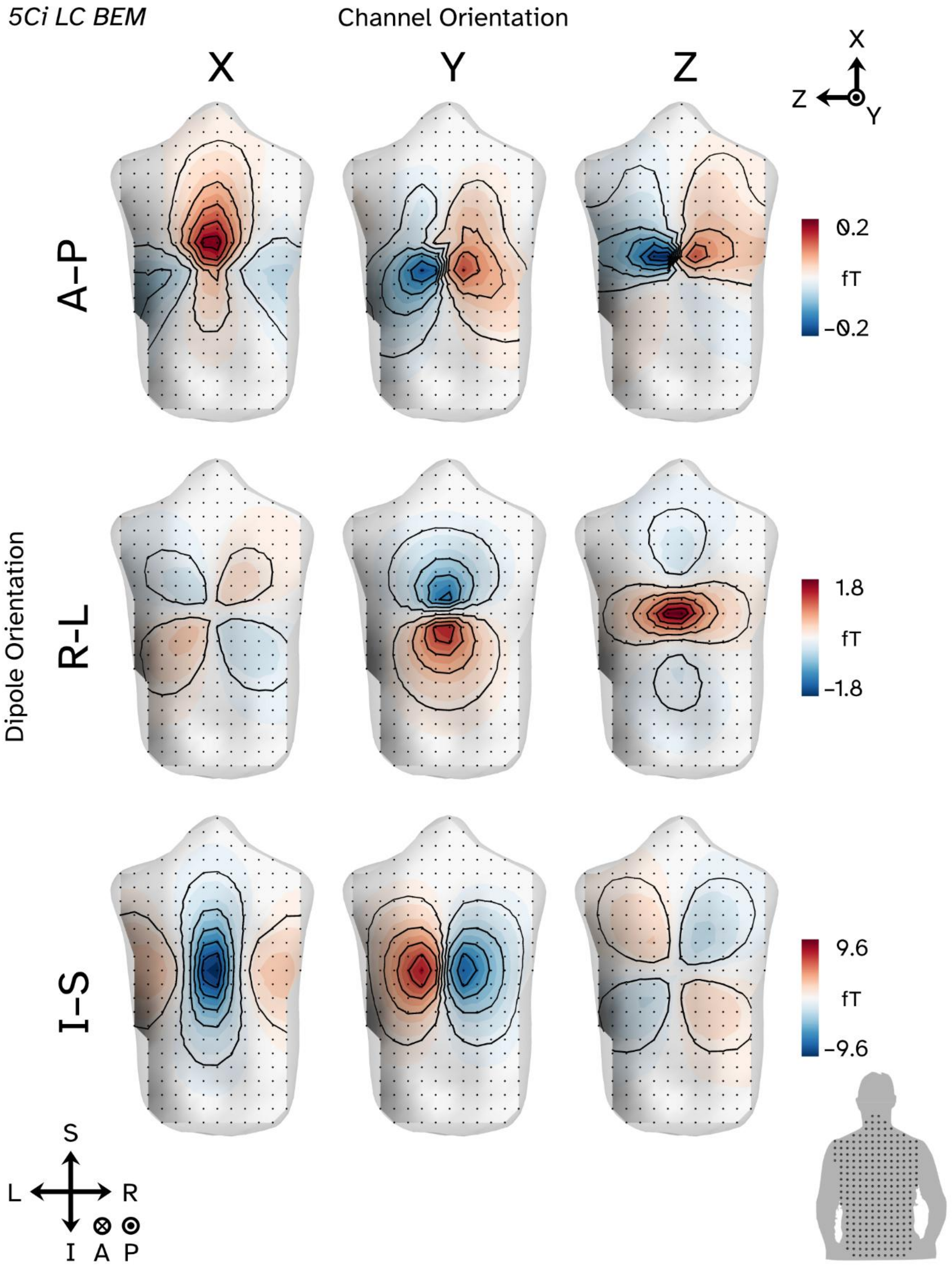
5Ci LC BEM
Channel Orientation
X
Y
Z
X
Z
Y
A–P
R–L
I–S
Dipole Orientation
0.2
fT
−0.2
1.8
fT
−1.8
9.6
fT
−9.6
S
L
R
I
A
P

Figure S13

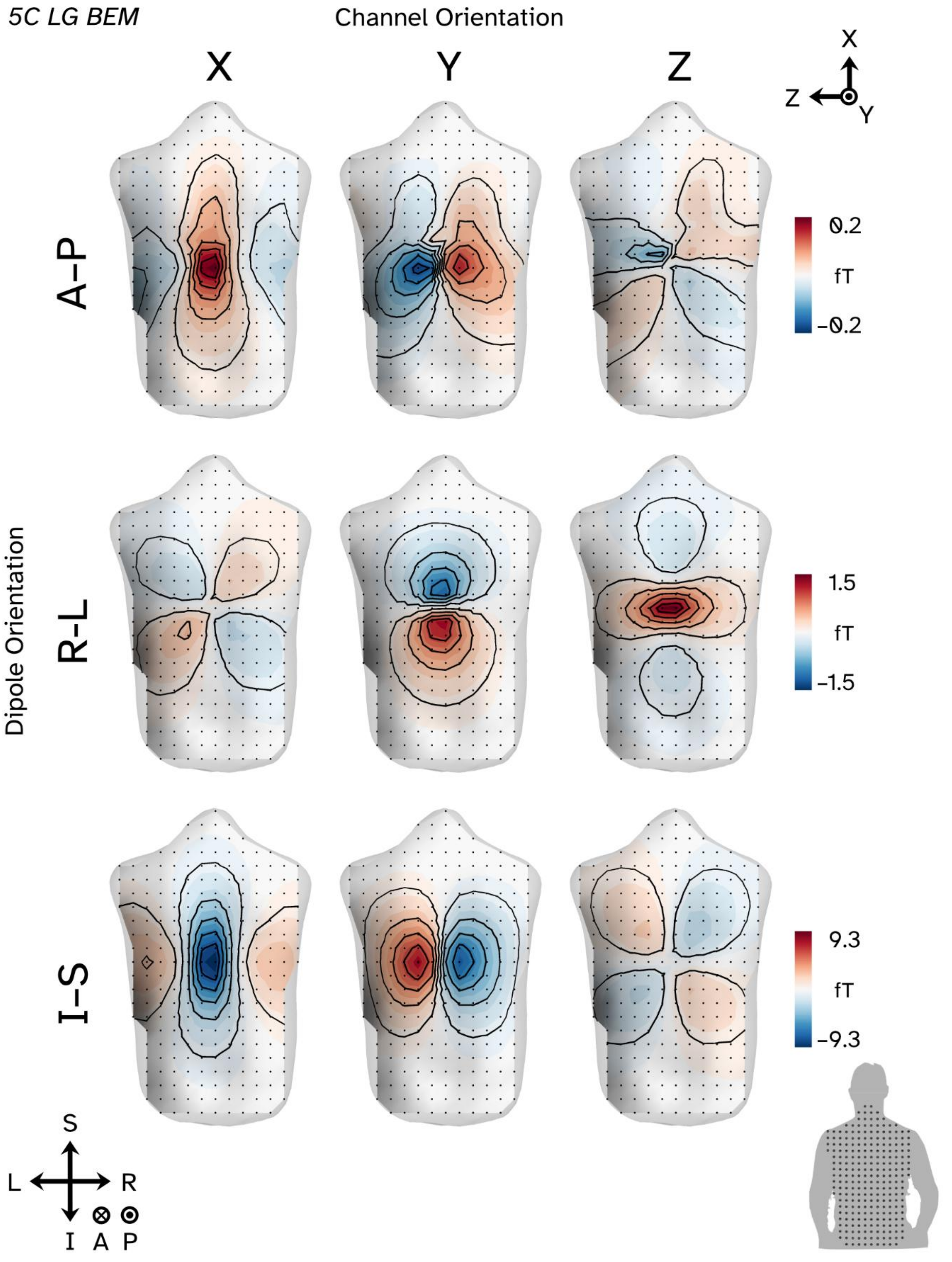
5C LG BEM
Channel Orientation
X
Y
Z
X
Z
Y
A–P
R–L
I–S
Dipole Orientation
0.2
fT
−0.2
1.5
fT
−1.5
9.3
fT
−9.3
S
L
R
I
A
P

Figure S14

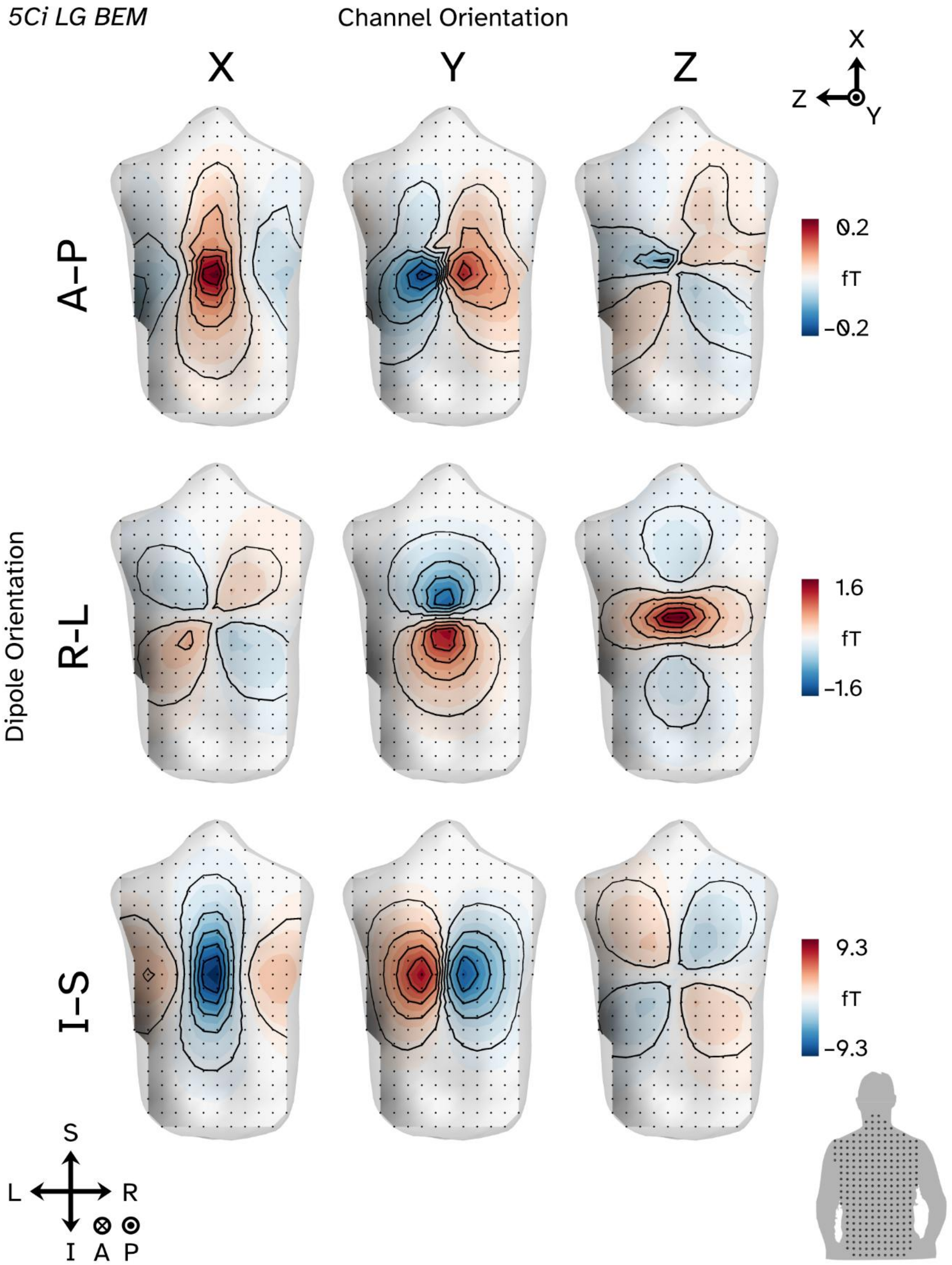
5Ci LG BEM
Channel Orientation
X
Y
Z
X
Z
Y
Dipole Orientation
A–P
R–L
I–S
0.2
fT
−0.2
1.6
fT
−1.6
9.3
fT
−9.3
S
L
R
I
A
P

Figure S15

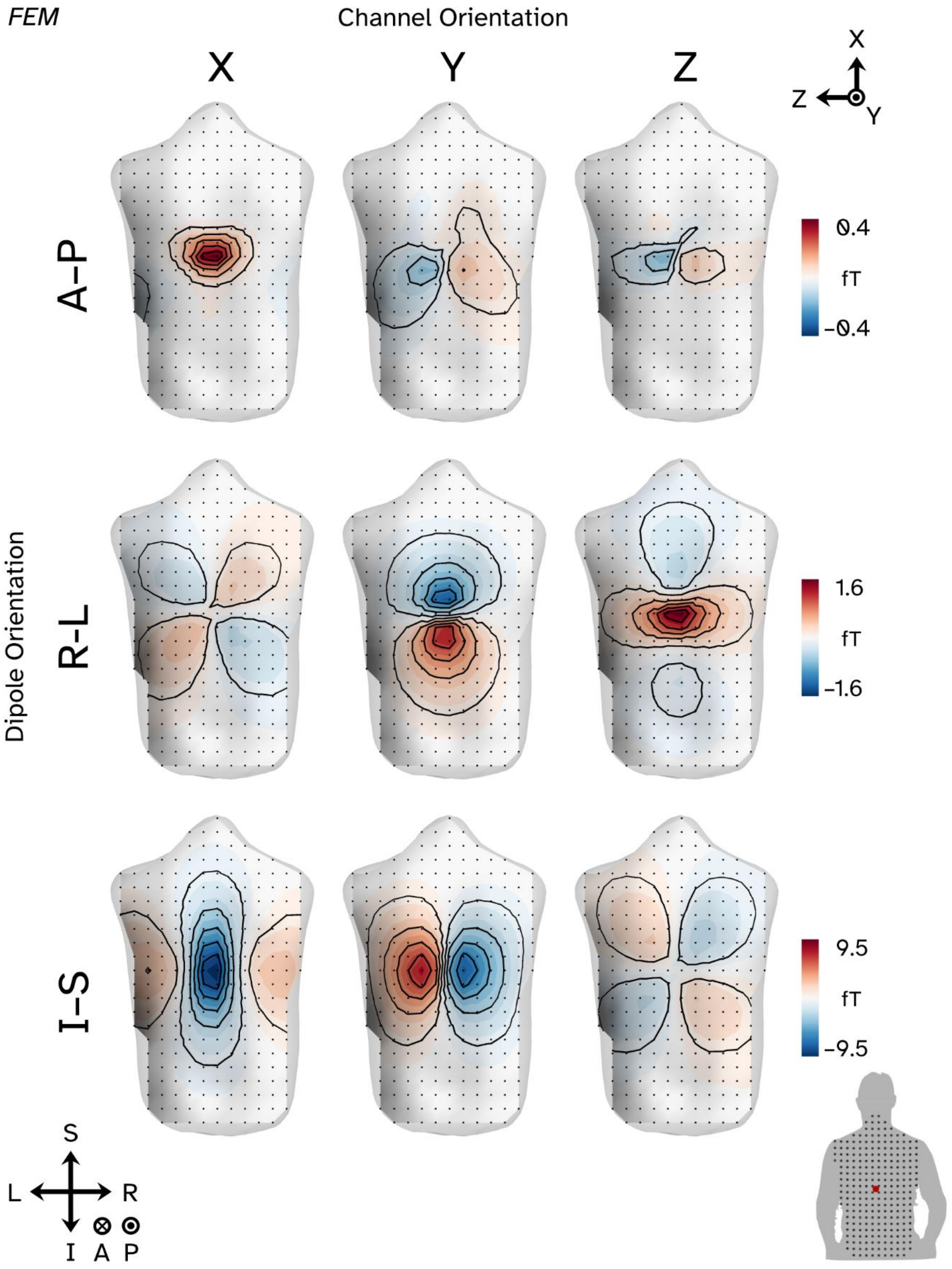
FEM
Channel Orientation
X
Y
Z
X
Z
Y
Dipole Orientation
A–P
R–L
I–S
0.4
fT
−0.4
1.6
fT
−1.6
9.5
fT
−9.5
S
L
R
I
A
P

Figure S16
